# Towards genetic indicators in ectomycorrhizal fungi: estimating the effective population size

**DOI:** 10.64898/2026.06.30.735680

**Authors:** Anouck Champion, Anna Bazzicalupo, Myriam Heuertz, Roberta Gargiulo

**Author notes:** Corresponding author: Roberta Gargiulo. These authors contributed equally to this work (Myriam Heuertz, Roberta Gargiulo).

## Abstract

Ectomycorrhizal (EM) fungi are vital to forest ecosystems, supporting tree growth and survival. However, their inclusion in conservation policy and action remains limited and little is known about the status of their genetic diversity, which is essential for their long-term survival and adaptation. The Global Biodiversity Framework adopted a genetic indicator based on the effective population size, *N*_e_, to monitor genetic diversity in all species. To date, it is still uncertain how *N*_e_, a key parameter, can be reliably assessed in species with complex life history traits. Ectomycorrhizal fungi are a highly diverse group of taxa displaying haplodiplontic life cycles with partially clonal reproduction. Here, we review the literature to understand how these life history traits might affect *N*_e_ and its estimation in six species of EM fungi. We estimated *N*_e_ in 19 populations using eight genetic and genomic datasets from selected studies. We compared *N*_e_ estimates using Linkage Disequilibrium (LD) and Sibship Frequency (SF) methods. We tested how *N*_e_ estimates change due to partial clonality and genetic structure gradients and whether the number of genetic markers influence the precision of the estimates. We show a systematic bias in *N*_e_ estimations when large clones are present and when populations are not correctly delimited. We found both methods are not robust to these factors, which makes them unreliable for conservation assessment purposes in EM fungi. This study provides new perspectives for further research into the links between life history traits and the effective population size of ectomycorrhizal fungi.

## Introduction

Genetic diversity is the basis for evolutionary changes and is critical for species’ long-term persistence and adaptation to changing environments (Hoban *et al*. 2020). However, genetic diversity is declining at an alarming pace because of habitat destruction and fragmentation, over-exploitation and other human activities, which significantly reduce population sizes (Hoban *et al*. 2021a, 2023). In recent years there has been remarkable progress in integrating the protection of genetic diversity in conservation policy, notably in the Convention on Biological Diversity’s (CBD) Kunming-Montreal Global Biodiversity Framework (GBF), adopted in 2022 (Conference of the parties to the convention on biological diversity 2022). The headline indicator A.4 of the GBF is the proportion of populations within species with an effective population size (*N*_e_) above 500, also known as the *Ne500* indicator, and was developed to assess and monitor genetic diversity in all species (Hoban *et al*. 2020). The effective population size is a genetic parameter determining the expected rate of genetic diversity loss due to genetic drift (Waples 2022). Empirical studies have shown that, when *N*_e_ < 500, genetic diversity is too low for a population to adapt, as random allele frequency variations due to genetic drift override directional changes due to natural selection (Jamieson & Allendorf 2012). Although this indicator has been shown to be relevant for a wide range of taxa (Hoban *et al*. 2021b; Mastretta-Yanes *et al*. 2024), challenges remain in estimating *N*_e_ in species where the definition of populations is complex (Fedorca *et al*. 2024). A major challenge is the accurate assessment of population delimitation, metapopulation dynamics and the level of connectivity between populations, which all affect *N*_e_ (Neel *et al*. 2013; Nunney 2016; Ryman *et al*. 2019; Waples & England 2011). This difficulty is compounded by taxon-specific life history traits, such as overlapping generations, mating systems and reproductive strategies that can affect *N*_e_ in contrasting directions (Fedorca *et al*. 2024; Waples 2005, 2016). Ectomycorrhizal fungi have several of these characteristics, which makes estimating their effective population sizes a complex task.

Mycorrhizal fungi are essential to terrestrial ecosystems, as many plant species depend on these symbionts for growth and survival (van der Heijden *et al*. 2015). Ectomycorrhizal (EM) fungi represent around 20,000 species forming diverse underground communities associated with around 6,000 plants (van der Heijden *et al*. 2015), including temperate, boreal and some tropical forest trees. Their complex life cycle alternates sexual reproduction with spore dispersal and asexual (clonal) propagation, spanning different ploidy states (haploid, dikaryotic, diploid), as shown in Figure 1. These features make it difficult to define individuals and populations, which hampers population genetic studies, and the estimation of *N*_e_ in particular (Dahlberg & Mueller 2011; Vincenot & Selosse 2017). Separate fruitbodies (e.g., mushrooms) are not necessarily different individuals because each mycelial genet (i.e. genetic individual) can produce several fruitbodies, or none at all (Vincenot & Selosse 2017). The effects of asexual reproduction on population genetic parameter estimation, including *N*_e_, have been explored in partially clonal plants (Gargiulo *et al*. 2023; Stoeckel *et al*. 2021b), but more research is required in ectomycorrhizal fungi, where the spatial and temporal dynamics of underground genets remain unclear (Vincenot & Selosse 2017). While population genetic theory assumes fully haploid or diploid organisms with random sexual reproduction, EM fungi depart from these characteristics. EM fungi are haplodiplontic species: organisms in which somatic development (here, mycelium) occurs both in the haploid and the diploid stages (Krueger-Hadfield *et al*. 2021). Haplodiplonty is widespread in the tree of life, occurring in algae, bryophytes, ferns, fungi, and appears to be a remarkably stable life cycle on evolutionary time scales (Stoeckel *et al*. 2021a). In addition, EM fungi populations are structured depending on how the spores are dispersed (by wind, soil mesofauna, mammals) and on the landscape. In areas devoid of barriers, EM fungi populations may display extensive gene flow, sometimes over hundreds of kilometres (Vincenot & Selosse 2017).

**Figure 1.**
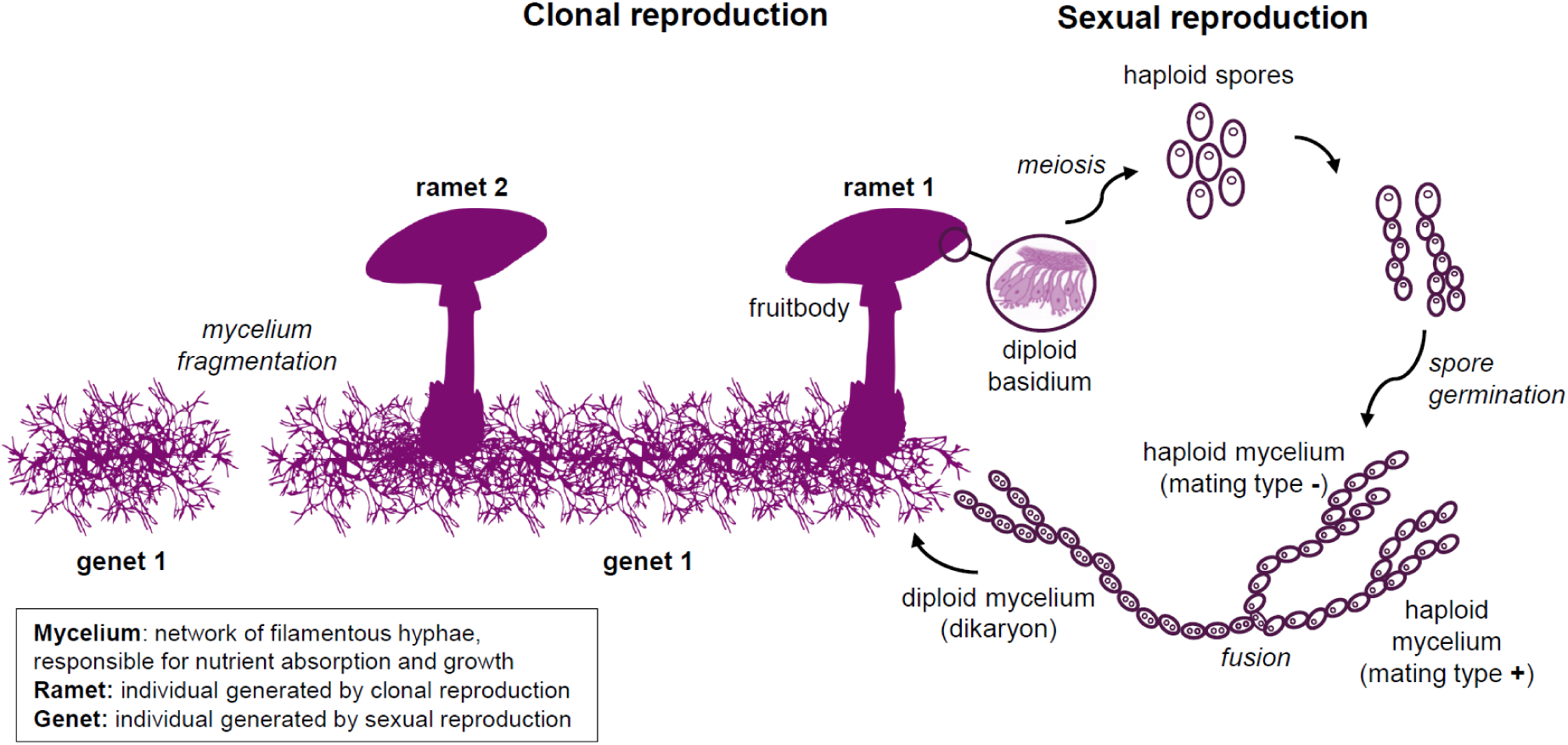
View of the typical life cycle of a basidiomycete EM fungi.

Recent population genetics and genomics studies based on fruitbody sampling have significantly advanced our understanding of EM fungi life cycles (Selosse *et al*. 2017), their adaptation to extreme environments including saline environments and metal-contaminated soils (Bazzicalupo *et al*. 2020; Branco *et al*. 2015) and have revealed genetic bases of local adaptation (Dauphin & Peter 2024). However, due to the difficulty in applying population genetic methods, conservation genetic studies are still rare for EM fungi and little is known about the loss of genetic diversity in these organisms (Abe *et al*. 2024). The data gathered in previous studies are valuable for continuing to assess the state of genetic diversity and explore population genetic processes in EM fungi. In order to apply the *Ne500* indicator, estimates of contemporary *N*_e_ are needed, which has seldom been assessed in EM fungi (Abe *et al*. 2024), although there are estimates of historical *N*_e_ variations across glacial oscillations (Tremble *et al*. 2023).

In practice, contemporary *N*_e_ can be estimated through several approaches: directly from demographic data (incorporating parameters such as sex ratio and variance of reproductive success), genetic data or, - when detailed data are unavailable - by approximation from the census size (*N*_c_, i.e., number of reproductively mature individuals) assuming *N*_e_/*N*_c_ = 0.1 (Laikre *et al*. 2021; Waples 2024b). For EM fungi, genetic methods are likely the most appropriate since demographic data are both sparse and challenging to collect. Contemporary *N*_e_ can be estimated from genetic data such as SNPs or microsatellites using single-sample methods, based on individuals sampled at a single time point (Do *et al*. 2014), or temporal methods when repeated samples are available. The two main single-sample methods are the linkage disequilibrium (LD) method (Waples 2024a) and the sibship frequency (SF) method (Wang 2009). The LD method is based on the estimation of linkage disequilibrium, i.e. the amount of non-random association of alleles at two independent loci (Waples 2024a), which is inversely proportional to *N*_e_ (Waples *et al*. 2016; Waples 2024a). This method assumes random mating, isolated populations and discrete generations, and many potential biases can arise when these assumptions are not respected (Waples 2024a). The SF method (Wang 2009) calculates the probability that two individuals taken at random in a sample are half or full-sibs, with *N*_e_ increasing as the relatedness between individuals decreases. The method assumes a closed population and discrete generations and is relatively robust to a small amount of immigration or departure from random mating (Wang 2009, 2016). Both methods require sufficient sample sizes and numbers of markers to obtain unbiased estimates, especially when *N*_e_ is large (Wang 2009; Waples 2024a), but this requirement is more pronounced for the SF method which becomes inaccurate unless the sample size is > 10% of the true *N*_e_ (Wang 2009).

The main objective of this study was to evaluate existing tools for the estimation of *N*_e_ in EM fungi in order to assess whether *N*_e_ is a feasible metric to monitor their genetic conservation status. Based on the reanalysis of published datasets, we estimated contemporary *N*_e_ in EM fungi species using both the LD and the SF methods. After identifying the theoretical and methodological challenges associated with *N*_e_ estimation in EM fungi (Table 1), we tested several hypotheses associated with each of the factors potentially affecting the reliability of the estimates. First, we investigated the effects of partial clonality on *N*_e_ estimation. Based on simulation studies, it is expected that populations with high rates of clonality (i.e. clonal reproduction > sexual reproduction) show higher values of *N*_e_ than strictly sexual populations, reflecting the fact that polymorphism is protected within individuals (Balloux *et al*. 2003). However, clonal reproduction also leads to a reduction in the number of different genotypes and an increase in LD, leading to a strong underestimation of *N*_e_ when using the LD-method (Gargiulo *et al*. 2023). Our aim was therefore to disentangle the impact of clonal reproduction on *N*_e_ and its estimation in EM fungi using empirical genotyping data and simulated sampling approaches. Second, we assessed the effect of population genetic structure on *N*_e_ estimates in EM fungi. Genotypic samples do not necessarily reflect homogeneous populations, and patterns of differentiation and migration can cause downwardly or upwardly biased *N*_e_ estimations if they are not accounted for, especially when using the LD-method (Waples 2024a; Waples & England 2011). Finally, we examined potential biases arising from sampling design in empirical genotyping studies of EM fungi, with particular focus on small sample sizes and missing data. We paid particular attention to issues associated with large genomic datasets (i.e. with large numbers of SNPs), where physical linkage between loci can lead to a downward bias in *N*_e_ estimation with the LD method (Waples 2024a). Datasets with a large number of SNPs also provide correlated information due to overlapping pairs of the same loci, which introduces pseudoreplication, affecting the precision and accuracy of *N*_e_ estimation (Waples *et al*. 2022).

**Table 1.** Main factors considered in this study, with related questions and expectations. The factors and questions marked by * have been explored in our article.

| Factor | Questions | Expectations & Hypotheses (reviewed from the literature) |
| --- | --- | --- |
| **Life-history traits** |  |  |
| **Partial clonality\*** | <u>Biological question</u><br>*What is the influence of clonality on the effective population size?*<br>*Do partially clonal species have a larger or smaller $N_e$ compared to strictly sexual species?* | - In partially clonal populations, asexuality is expected to increase heterozygosity, due to the absence of chromosome segregation and recombination (Ellegren & Galtier 2016). Hence, $N_e$ is expected to be high when the rate of clonality ($c$) is strong, polymorphism being fixed in clonal lineages. However, $N_e$ should not be significantly affected by clonality unless sexual reproduction becomes very rare ($c = 1$), where infinite values of $N_e$ are expected (Balloux *et al.* 2003; Stoeckel *et al.* 2021b).<br>- Variance in reproductive success is also expected to increase with higher clonal rates, which would result in a decreased $N_e$ (Balloux *et al.* 2003; Stoeckel *et al.* 2021a; Waples 2024b, 2025)<br>- When using coalescent based methods, $N_e/N_c$ ratios in organisms with clonal reproduction is lower compared to ratios measured in strictly sexual organisms. This is due to longer generation times occurring in clonal organisms, but does not necessarily imply low genetic diversity (the role of somatic mutations remains unclear). Furthermore, these lower ratios may also be caused by larger census size (Orive 1993). |
|  | <u>Methodological issue*</u><br>*Which estimation method is more robust to violation of non-* | - A downward bias is expected in the estimation of $N_e$ with the LD method in partially clonal populations. First, clonal reproduction results in replicated genotypes in the datasets. This situation breaks the assumption of random mating, to which the LD-method is very sensitive (Waples 2024a). Moreover, clonal populations display more LD due to the absence of recombination (Stoeckel *et al.* 2021b). |
|  | *random mating associated with clonal reproduction?* | - The SF method is not particularly sensitive to deviations from random mating, thus we expect this method to be less affected by the presence of replicated genotypes (Wang 2009). |
| **Haplo-diplontic life-cycle** | <u>Biological question</u><br>*What is the combined effect of haplodiplonty and partial clonality on the effective size of EM fungi ?* | - As in diploid and haploid organisms, genotypic diversity is expected to decrease with increasing clonality. However, the number of haploids constrains the number of possible genotypes. Thus, for high haploid population sizes, the number of possible combinations of haplotypes is increased, leading to a higher genotypic richness (Krueger-Hadfield *et al.* 2021).<br>- Contrarily, simulations show that maximum values of $N_e$ are reached for lower proportions of haploids and increased levels of clonality (Stoeckel *et al.* 2021a). |
|  | <u>Methodological issue</u><br>*How can haploid and diploid populations of EM fungi be compared simultaneously?* | - As recommended in Krueger-Hadfield *et al.* (Krueger-Hadfield *et al.* 2021), both the haploid and the diploid pools must be studied when looking at haplodiplontic species. However, spore haplotypes are difficult to obtain in EM fungi, and only diploid information is available for most species.<br>- Moreover, the size of the haploid compartment is unknown, due to the difficulty to isolate and genotype haploid mycelium. Thus, even if spore production is very high, the proportion of spores surviving to form the haploid pool is difficult to assess. |
| **Population structure** |  |  |
| **Genetic structure\*** | <u>Biological question</u> | - In EM fungi species that reproduce mainly sexually, the wind-dispersion of spores would result in high population connectivity, with patterns of isolation by distance. On the contrary, species reproducing sexually with underground fruitbodies (dispersed by animals) might display more discontinuous populations (Vincenot & Selosse 2017). |
|  | *What is the effect of genetic structure on $N_e$ in ectomycorrhizal fungi?* | - Different genetic and spatial genetic structures are likely to affect $N_e$ (Nunney 2016). |
|  | <u>Methodological issue*</u><br>*Are current methods for estimating $N_e$ biased by the genetic structure of ectomycorrhizal fungi populations?* | - The LD-method assumes isolated populations, but this assumption is rarely met in natural populations. Thus, two biases are expected: (i) a downward bias due to Wahlund effect (increase of mixture LD), and (ii) an upward bias due to migration which increases the number of potential parents (decreases the drift LD) (Waples 2024a; Waples & England 2011).<br>- As long as there is no direct immigrant in the sample, the SF method is not affected by the presence of descendents of immigrants in the sample (Wang 2009).<br>- However, single-sample, contemporary estimators (such as LD and SF estimators) are likely to be very sensitive to the size of the local breeding window of the species (Neel *et al.* 2013). |
| **Sampling design** |  |  |
| **Number of markers\*** | <u>Methodological issue*</u><br>*What is the effect of large genomic datasets on precision when using LD or SF based methods for $N_e$ estimation?* | - When using large genomic datasets (e.g. thousands of SNPs), some markers are likely to be physically linked and/or provide correlated information.<br>- When using the LD-method, precision is reduced due to overlapping pairs of the same loci, when using more than a few thousand loci. This effect is expected to be even stronger in populations with small $N_e$ (pedigree-dependence). Large confidence intervals should be expected, while common methods (jackknife) often overestimate precision (narrow CIs), compared with a situation where the loci are independent (Waples *et al.* 2022). |
| **Sample size\*** | <u>Methodological issue*</u><br>*What sample size is required to obtain an accurate estimate of $N_e$?* | - When using genomic-scale datasets, both SF and LD methods provide unbiased estimates from rather small sample sizes ($n \sim 30$), as long as the true $N_e$ is not too large ($N_e$ 50-5,000) (Waples 2021). This is because, in small populations, genetic drift is strong, thus $N_e$ is easy to estimate (Wang 2025).<br>- However, for very large populations ($N_e > 10,000$), both methods are inaccurate unless the sample size is large enough, the LD method being more biased, but also more precise, than the SF method (Wang 2025). |

Our re-analysis of population genetic datasets enabled us to unravel the factors that might affect the reliability of *N*_e_ estimates and to suggest ways of mitigating their effect for improved estimations of *N*_e_ and, more generally, genetic diversity in EM fungi.

## Material and Methods

### Ectomycorrhizal fungi genotypic datasets

#### Selection of datasets

To find genotypic datasets, a keyword search in NCBI was carried out using the following query: mushroom* AND (“microsatellites” OR “SNPs”) AND *mycorrhiza*. The term “mushroom” was preferred to “fungi” in order to exclude fungi that do not form fruitbodies (“mushrooms” in the general sense), as most genetic studies on EM fungi are based on fruitbodies sampling. A first filter was applied to keep only population genetic studies on EM fungi (removing publications about ericoid or orchid mycorrhizal fungi). Then the search was limited to microsatellite and SNPs markers. Additional datasets were found via review articles or by contacting authors directly, until the end of 2024. Only datasets from Basidiomycetes species were selected, as most Ascomycetes datasets (e.g., from the well-known species *Tuber* spp.) represent haploid genotypes, and, to date, no software programmes support this kind of data for *N*_e_ estimation.

#### Description of datasets

Eight genetic datasets from six EM species were used in this study: *Boletus edulis* Bull. (Brejon Lamartinière *et al*. 2024; Hoffman *et al*. 2020), *Suillus brevipes* (Peck) Kuntze (Branco *et al*. 2017), *Suillus luteus* (L.) Roussel (Bazzicalupo *et al*. 2020; Pildain *et al*. 2021), *Laccaria amethystina* Cooke (Vincenot *et al*. 2012), *Rhizopogon togasawarius* Mujic et al. (Abe *et al*. 2024) and *Cantharellus formosus* Corner (Dunham *et al*. 2006). These datasets include species ranging from cosmopolitan (*L. amethystina*, *B. edulis*) to narrowly endemic (*R. togasawarius*), with populations sampled from both native and recently introduced ranges (Figure S1). The main characteristics of these datasets are presented in Table 2.

**Table 2.** Characteristics of the eight genotypic datasets used in this study. For each dataset, the total sample size is indicated in the fourth column. If the datasets contained replicated genotypes, the number of corresponding Multi-Locus Genotypes (MLGs) is indicated. From the original datasets, some samples and/or populations were not used; the populations considered in this study are indicated in the “Populations” column, with corresponding sample sizes.

| Species name | Taxonomy | Ecology | Total sample size | Total number of markers | Populations (sample sizes) | Issues explored | Reference |
| --- | --- | --- | --- | --- | --- | --- | --- |
| *Boletus edulis*<br>Bull. | Basidiomycota ><br>Boletales ><br>Boletaceae | Generalist, associated with pine, birch, beech and oak species. Widespread across Eurasia and North America. | 134<br>(43 MLGs) | 7 SSRs | Bielefeld_1 (19)<br>Bielefeld_2 (22) | Influence of partial clonality | (Hoffman *et al.* 2020) |
| *Boletus edulis*<br>Bull. |  |  | 110 | ~ 21 M SNPs | Europe (49)<br>West Coast (35)<br>East Coast (26) | Influence of pseudoreplication | (Brejon Lamartinière *et al.* 2024) |
| *Suillus brevipes*<br>(Peck) Kuntze | Basidiomycota ><br>Boletales ><br>Suillaceae | Obligate symbiont of *Pinus* species. Widespread in North America. | 55 | 662940 SNPs | California (25)<br>Minnesota/Alberta (18)<br>Wyoming/Colorado (12) | Influence of genetic structure, influence of pseudoreplication | (Branco *et al.* 2017) |
| *Suillus luteus*<br>(L.) Roussel | Basidiomycota ><br>Boletales ><br>Suillaceae | Associated with two-needle pines. Native to the Northern hemisphere, introduced in South America, South Africa and Oceania. | 34 | ~ 1.6 M SNPs | Belgium (34) | Influence of pseudoreplication | (Bazzicalupo *et al.* 2020) |
| *Suillus luteus*<br>(L.) Roussel |  |  | 106 | 5 SSRs | Patagonia (106) | Influence of partial clonality | (Pildain *et al.* 2021) |
| *Laccaria amethystina*<br>Cooke | Basidiomycota ><br>Agaricales ><br>Hydnangiaceae | Generalist, found in most Northern temperate zones, both in coniferous and deciduous forests. | 632 | 9 SSRs | Europe (564)<br>Japan - Mount Fuji (36)<br>Japan - Mount Tokachi (32) | Influence of partial clonality, influence of genetic structure | (Vincenot *et al.* 2012) |
| *Rhizopogon togasawarius*<br>Mujic *et al.* | Basidiomycota ><br>Boletales ><br>Rhizopogonaceae | Specific to *Pseudotsuga japonica*. Endemic and endangered species in Japan. | 236 | 20 SSRs | Kawamatakannon - KW (53)<br>Sannokogawa - SN (44)<br>Ohmata - OM (18)<br>Nishinoko - NK (86)<br>Yasudagoyama - YS (35) | Influence of partial clonality, influence of genetic structure | (Abe *et al.* 2024) |
| *Cantharellus formosus*<br>Corner | Basidiomycota ><br>Cantharellales ><br>Cantharellaceae | Native to the Pacific Northwest region of North America. Abundant in coniferous forests. | 285<br>(19 MLGs) | 5 SSRs | Oregon (19) | Influence of partial clonality | (Dunham *et al.* 2006) |

### Data processing and filtering

Most analyses were conducted in R 4.3.2 (R Core Team 2021), using the Rstudio interface, and on the Genotoul bioinformatics platform. Starting from the original data provided by the authors, loci with more than 20% of missing data across all individuals were excluded systematically using the R package *poppr* (Kamvar *et al*. 2014) for SSR data, and using the “max-missing” option in VCFtools for SNP data. Genotypes with more than 20% missing data across all loci were also removed.

#### Genetic structure analysis and genetic diversity indices

As most *N*_e_ estimation methods assume isolated populations (especially the LD-method), a genetic structure analysis was carried out on each dataset in order to identify discrete genetic clusters (i.e. putative populations). Physically linked SNPs were removed with the software programme PLINK2 (Chang *et al*. 2015), using the “indep-pairwise” option, to meet the assumption of non-linked markers required by STRUCTURE. Parameters were set to 50 and 0.5 for the window size and r2 threshold, respectively. For the SSRs datasets, we assumed that the original studies developing the markers had carried out a routine test for genotypic disequilibrium, and therefore that the SSR loci used were unlinked. All datasets were clone-corrected before genetic structure analysis, to minimise the influence of clonality on genetic structure. The R packages *poppr* (Kamvar *et al*. 2014) and *Rclone* were used to check for the presence of replicated genotypes in SSR datasets, and to keep only one copy per MLG (Multi-Locus Genotype). For SNPs dataset, replicated genotypes had already been removed by the authors. Genetic structure was assessed using the programme STRUCTURE 2.3.4 (Pritchard *et al*. 2000), using a subset of 20,000 SNPs for the genomic datasets, and the whole datasets for microsatellite datasets. Outputs from STRUCTURE were analysed with the online programme StructureSelector (Li & Liu 2018). The best number of clusters (K) was determined by comparing several indicators in StructureSelector, and the interpretation of barplots allowed to assign each individual to a cluster (Figures S2 to S9). Each of these clusters were then considered as a discrete population for further analysis (see ‘Populations’ in Table 2). For the datasets of *Rhizopogon togasawarius* (Abe *et al*. 2024) and *Suillus luteus* (Bazzicalupo *et al*. 2020; Pildain *et al*. 2021), the corresponding publications already contained the STRUCTURE results and the analysis was not repeated.

Measures of genetic diversity were calculated for each clone-corrected population using the R package *hierfstat* (Goudet & Jombart 2022) for the microsatellite datasets: Allelic richness (*A*_r_, mean across all loci), observed heterozygosity (*H*_o_), expected heterozygosity (*H*_E_), inbreeding coefficient (*F*_IS_) and its significance. The same summary statistics and nucleotide diversity (π) were computed for the SNPs datasets using the *dartR* (Gruber *et al*. 2018) and *adegenet* (Jombart & Ahmed 2011) packages.

### *N*_e_ estimation

#### Estimation of effective population size

We computed a first estimation of *N*_e_ using the methods described below for each population, using clone-corrected datasets and a subset of 2,000 SNPs for the genomic datasets, in order to obtain comparable estimates across populations and species. Then, we explored the influence of partially clonal reproduction, genetic structure, and the number of markers on the genetic estimation of *N*_e_ (see main questions and hypotheses in Table 1). The estimation of the contemporary *N*_e_ was performed using two different methods: the linkage disequilibrium method (LD), using the software NeEstimator v2 (Do *et al*. 2014) and the sibship frequency method (SF) using the program Colony2 v2.0.7.1 (Jones & Wang 2010).

The sibship frequency method developed by Wang (2009) calculates the probability that two individuals taken at random in a sample are half or full-sibs, which allows to estimate the *N*_e_ of the population. In Colony, we used the Full-Likelihood (FL) method for sibship and parentage assignment with the following settings: “Female Polygamy”, “Male Polygamy”, “Without inbreeding”, “Without clone”, “Dioecious”, “Diploid”. All other parameters were kept to default values. An analytical pipeline was created to format data from VCF or GENEPOP files to Colony-like input files and run multiple analyses in Colony from a computing cluster, thereby significantly improving file processing speed compared to the GUI version of Colony. The program provides two estimates of *N*_e_: a ‘random-mating’ estimation, where the population is assumed to be at Hardy-Weinberg equilibrium, and a ‘non-random mating’ estimation, adjusted using the calculation of alpha (the deviation from Hardy-Weinberg equilibrium) from the genotype data.

The LD method relies on the measure of linkage disequilibrium between pairs of loci to estimate *N*_e_. In NeEstimator, we used the criterion “random mating” and the “no singleton alleles” critical value for allele frequencies. Removing singleton alleles has been proposed as the best way to avoid bias related to rare alleles, while preserving valuable demographic information (Waples 2024a). The NeEstimator programme was used with the R interface provided by the *RLDNe* package (Robinson 2019).

#### Effect of partial clonality on N_e_ estimation

Both the LD and the SF methods are known to be differently influenced by non-random mating, and partially clonal reproduction is precisely a case of deviation from a random mating system. To assess how *N*_e_ estimations are affected by the presence of replicated genotypes in a dataset, we first used datasets in which clonality has been observed: *Boletus edulis* (Hoffman *et al*. 2020) and *Cantharellus formosus* (Dunham *et al*. 2006). These two datasets included replicated genotypes (i.e. several ramets from the same genet), and thus can provide an initial estimate of the level of clonality in these species. Clonal rates (c) in each population were assessed from the original datasets (with replicated genotypes), using genetic and genotypic indices (Stoeckel *et al*. 2021b) in the R package *Rclone* (Arnaud-Haond & Bailleul 2022): Genotypic richness (R), Pareto’s beta, Inbreeding coefficient (F_IS_) and linkage disequilibrium (rd). However, these estimates of clonal rates remain highly uncertain due to the sampling approach adopted in the original studies. For instance, clones might have been excluded from sampling, as in Dunham *et al*. (2006), where fruitbodies were sampled at a minimum distance of 5 m between two fruitbodies; and some genets might not be represented by fruitbodies for several years (only one year of sampling in both cases).

To reduce our uncertainty regarding clonality, we artificially manipulated clonality rates by removing or adding replicated genotypes in one population in each of the five following species: *Boletus edulis* (Hoffman *et al*. 2020), *Cantharellus formosus* (Dunham *et al*. 2006), *Suillus luteus* (Pildain *et al*. 2021), *Laccaria amethystina* (Vincenot *et al*. 2012) and *Rhizopogon togasawarius* (Abe *et al*. 2024). The aim was to obtain gradients of clonality for several species, and verify how *N*_e_ estimates varied across these gradients. For each population, the presence of clonal individuals was assessed using the R packages *poppr* (Kamvar *et al*. 2014) and *Rclone* (Arnaud-Haond & Bailleul 2022). One copy of each MLG (Multi-Locus Genotype, i.e. identical genotype) was kept in each dataset, and these clone-corrected datasets were considered the “original without clones” datasets. From each original dataset, we constructed a clonality gradient by creating 4 different conditions. (1) “Hardy-Weinberg genotypes” (HW): genotypes were drawn randomly from the clone-corrected original dataset, based on observed allelic frequencies. This represented a population following Hardy-Weinberg proportions, and appeared as the lowest clonality in our gradient. (2) “Original without clones” (NC): the clone-corrected original dataset. (3) “Few clones” (FC): each original genotype is replicated 2-4 times to increase clonality. (4) “Large clone” (LC): one genotype is drawn from the population and replicated 10-20 times to represent a large clonal lineage. A resampling corresponding to the sample size of the original dataset was performed in each condition to obtain comparable sample sizes across the gradient. MLG plots of all four conditions in five species are shown in Figure S10. Finally, *N*_e_ was estimated in each condition across the five species, using the LD and SF methods. Significant differences between the four datasets across the five populations were tested using a Kruskal-Wallis rank sum test, followed by a Dunn post hoc test (adjusted using the Holm method).

#### Effect of genetic structure on N_e_ estimation

The effect of genetic structure on *N*_e_ estimations was tested in populations of species with contrasted dispersal modes and spore production: *Suillus brevipes* (Branco *et al*. 2017), *Laccaria amethystina* (Vincenot *et al*. 2012) and *Rhizopogon togasawarius* (Abe *et al*. 2024). For each dataset, *N*_e_ was estimated at the population level (as defined in Table 2), at the regional level (by combining several populations) and at the subpopulation (sampling site, where relevant) level, to assess how population delimitation affects *N*_e_ estimations.

In *Rhizopogon togasawarius*, five populations were identified by the authors (NK, YS, KW, OM, SN, see Figure 1 in Abe *et al*. 2024), with strong pairwise *F*_ST_ values between sites. This level was retained as the population level. The STRUCTURE analysis indicated also a high probability of two-group clustering, corresponding to both islands isolated by a sea barrier. We retained this as the regional level (see Figure S11) for *N*_e_ estimation (Shikoku Island = NK + YS; Kii Peninsula = KW + OM + SN). In *Suillus brevipes*, the STRUCTURE analysis identified K=3 genetic clusters (California, Minnesota/Alberta, Wyoming/Colorado, see Figure S6) retained as the population level. We retained K=2 as the regional level, distinguishing California from the merged inland populations (Minnesota/Alberta + Wyoming/Colorado). In *Laccaria amethystina*, extensive gene flow has been reported across large geographical ranges, resulting in three main clusters: Europe, Japan-Mount Fuji and Japan-Mount Tokachi (situated on two different islands in Japan), here considered as the population level (see Table 2). For the European population, the substructure was also taken into account (subpopulations corresponding to sampling sites, see Figure S7) and Japanese samples were combined into a single cluster (Figure S8), representing a regional level.

In addition to *N*_e_ estimation at the subpopulation (subpop-*N*_e_), population (*N*_e_) and regional levels (regional-*N*_e_), the sum of estimates made at the population level (∑*N*_e_) were calculated, to compare it with the regional-*N*_e_, as in Mergeay *et al*. (2024). If no bias were at play, we would expect the regional-*N*_e_ to be equal to the sum of *N*_e_ estimates obtained at the population level.

#### Effect of number of markers on N_e_ estimation

Several studies have pointed out that the use of genomic-scale datasets for the estimation of *N*_e_ with the LD method can affect both the accuracy and the precision of the estimates (Waples 2016; Waples *et al*. 2022). Also, biases can be expected when using the SF method with large genomic datasets, as this method was designed for datasets with a relatively small number of markers (10–100 markers). On the contrary, SNP datasets often contain several thousand or even millions of SNPs, which raises the question of the applicability of this method to such datasets. In order to evaluate the accuracy and the precision of *N*_e_ estimation related to the use of large numbers of SNPs on both methods, we used three SNP datasets deriving from whole genome resequencing of *Suillus brevipes* (Branco *et al*. 2017), *Suillus luteus* (Bazzicalupo *et al*. 2020) and *Boletus edulis* (Brejon Lamartinière *et al*. 2024). From the original datasets containing around 662K, 1.6M and 21M SNPs for *Suillus brevipes*, *Suillus luteus* and *Boletus edulis* respectively, random subsets of 2,000, 5,000 and 10,000 SNPs were produced using VCFtools (Danecek *et al*. 2011), with 10 replicates for each population. *N*_e_ was then estimated for each replicate using the LD and SF methods. A Levene test was used to assess homogeneity of variances of *N*_e_ estimates across the SNPs gradient in each population, thus assessing accuracy of each method with various number of markers.

## Results

### Population delimitation and genetic indices

Following data processing, filtering and analysis of genetic structure (see Figures S2 to S9), our dataset included 19 genetic populations across six species of ectomycorrhizal fungi (Table 2). Genetic diversity metrics computed from clone-corrected datasets for each population are shown in Table 3. Allelic richness estimates for the five SSR datasets are homogenous across populations within each species, except in *Boletus edulis*, in which *A*_R_=3.4 alleles in Bielefeld_1 vs *A*_R_=6.3 in Bielefeld_2. Expected heterozygosity (*H*_E_, for SSR datasets) and nucleotide diversity (π, for SNP datasets) were also similar across populations within species for each marker type. Inbreeding coefficients (*F*_IS_) were significant in about half of the studied populations, with a stronger deficit of heterozygotes observed in populations that merged multiple sampling sites (e.g., European populations in *Boletus edulis* and *Laccaria amethystina*). Table 3 also includes *N*_e_ estimates with the Linkage Disequilibrium (LD) and Sibship Frequency (SF) methods for each population. *N*_e_ estimates obtained with the sibship frequency (SF) method in Colony typically lay below 100, except in the Europe population of *L. amethystina*. The “non-random mating” option yielded systematically lower estimates than the “random-mating” option, while both were of the same order of magnitude. The LD method also yielded *N*_e_ point estimates typically below 100, but with confidence intervals often extending to infinity, or even to negative estimates. Only two populations exceeded the 500 threshold defined for the *Ne500* indicator: the Belgium population of *Suillus luteus* and the Europe population of *Laccaria amethystina*.

**Table 3.** Genetic indices for each population. Abbreviations: N, sample size; *A*_R_, Allelic richness (only for SSR data); ***π***, Nucleotide diversity (for SNP data); *H*_E_, Expected Heterozygosity (for SSR data); *F*_IS_, Inbreeding coefficient; Signif, Result of bootstrapping test for *F*_IS_; SF-*N*_e_ (rm), Estimation of *N*_e_ with SF (random mating); SF-*N*_e_ (nrm), Estimation of *N*_e_ with SF method (non random mating); 95% CI, 95% confidence interval calculated from Student’s distribution; LD-*N*_e_, Estimation of *N*_e_ with LD method; Jackknife CI, Jackknife confidence interval.

| Population | N | Marker type | Nbr markers | $A_R$ | $\pi$ | $H_E$ | $F_{IS}$ | Signif | $SF-N_e$ (rm) | 95% CI | $SF-N_e$ (nrm) | 95 % CI | $LD-N_e$ | Jackknife CI |
| --- | --- | --- | --- | --- | --- | --- | --- | --- | --- | --- | --- | --- | --- | --- |
| ***Boletus edulis*** |  |  |  |  |  |  |  |  |  |  |  |  |  |  |
| Bielefeld 1 | 19 | SSR | 7 | 3.397 |  | 0.458 | -0.011 | ns | 10 | 5 - 26 | 7 | 3 - 21 | 1.4 | 0.6 - 3.9 |
| Bielefeld 2 | 22 | SSR | 7 | 6.285 |  | 0.638 | 0.168 | * | 14 | 7 - 31 | 7 | 4 - 21 | 23 | 3.2 - Inf |
| East Coast | 26 | SNP | 2000 |  | 0.057 |  | 0.061 | ns | 45 | 28 - 82 | 38 | 23 - 67 | 133.8 | 43.4 - Inf |
| Europe | 49 | SNP | 2000 |  | 0.047 |  | 0.174 | * | 39 | 25 - 62 | 31 | 19 - 53 | 93.9 | 35.6 - Inf |
| West Coast | 35 | SNP | 2000 |  | 0.037 |  | 0.054 | ns | 21 | 12 - 43 | 23 | 14 - 44 | 116.9 | 50.3 - Inf |
| ***Cantharellus formosus*** |  |  |  |  |  |  |  |  |  |  |  |  |  |  |
| Oregon | 19 | SSR | 5 | 3 |  | 0.395 | -0.093 | ns | 10 | 5 - 28 | 9 | 4 - 24 | Inf | 12.5 - Inf |
| ***Laccaria amethystina*** |  |  |  |  |  |  |  |  |  |  |  |  |  |  |
| Europe | 564 | SSR | 9 | 3.089 |  | 0.456 | 0.334 | * | 258 | 214 - 308 | 124 | 97 - 162 | 991.9 | 251.8 - Inf |
| Japan - Mount Fuji | 36 | SSR | 9 | 3.381 |  | 0.461 | 0.156 | ns | 19 | 11 - 38 | 11 | 6 - 30 | -82.4 | 59.8 - Inf |
| Japan - Mount Tokachi | 32 | SSR | 9 | 2.737 |  | 0.444 | 0.186 | ns | 12 | 6 - 26 | 8 | 4 - 23 | -39.7 | 53.5 - Inf |
| ***Rhizopogon togasawarius*** |  |  |  |  |  |  |  |  |  |  |  |  |  |  |
| Kawamatakannon | 53 | SSR | 20 | 2.943 |  | 0.368 | 0.12 | * | 37 | 24 - 61 | 24 | 14 - 43 | 65.1 | 29.1 - 459.5 |
| Nishinoko | 86 | SSR | 20 | 4.004 |  | 0.466 | 0.049 | * | 39 | 26 - 62 | 31 | 20 - 52 | 85.1 | 56 - 149.2 |
| Ohmata | 18 | SSR | 20 | 3.1 |  | 0.474 | 0.098 | ns | 16 | 8 - 36 | 10 | 5 - 28 | 23.5 | 2.2 - Inf |
| Sannokogawa | 44 | SSR | 20 | 3.989 |  | 0.442 | 0.106 | * | 28 | 17 - 48 | 18 | 11 - 35 | 41.6 | 24.6 - 87.5 |
| Yasudagoyama | 35 | SSR | 20 | 3.765 |  | 0.532 | 0.106 | * | 22 | 13 - 42 | 14 | 8 - 31 | 27.5 | 14.3 - 71.7 |
| ***Suillus brevipes*** |  |  |  |  |  |  |  |  |  |  |  |  |  |  |
| California | 25 | SNP | 2000 |  | 0.133 |  | 0.04 | ns | 52 | 31 - 95 | 52 | 30 - 98 | 48.9 | 30.3 - 93.6 |
| Minnesota / Alberta | 18 | SNP | 2000 |  | 0.167 |  | 0.137 | ns | 41 | 21 - 22 | 33 | 16 - 89 | 13.4 | 8.3 - 25.8 |
| Wyoming / Colorado | 12 | SNP | 2000 |  | 0.137 |  | 0.204 | * | 38 | 17 - 8 | 26 | 11 - 3 | 2.5 | 1.1 - 23.1 |
| ***Suillus luteus*** |  |  |  |  |  |  |  |  |  |  |  |  |  |  |
| Patagonia | 96 | SSR | 5 | 8.88 |  | 0.622 | 0.114 | ns | 43 | 29 - 67 | 25 | 15 - 43 | 214.9 | 38.8 - Inf |
| Belgium | 34 | SNP | 2000 |  | 0.185 |  | 0.041 | * | 86 | 50 - 97 | 68 | 39 - 58 | 3797.9 | 361.5 - Inf |

### Influence of simulated partial clonality on Ne estimation

Population genetic datasets contained variable numbers of replicated multi-locus genotypes (MLGs) though clone-correction by the authors in some studies precluded a comparative assessment across species. The highest number of ramets per genet was observed in *Cantharellus formosus* (Dunham *et al*. 2006) and *Boletus edulis* (Hoffman *et al*. 2020)(Figure S12). The *Cantharellus formosus* dataset contained one large clone with more than 120 ramets, and thus featured a low genotypic richness (R = 0.06) and a low Pareto’s beta value (0.2)(Figure S13). In *Boletus edulis*, most MLGs were represented by less than 20 ramets, which resulted in higher genotypic richness (R = 0.34) and a more even distribution of replicated genotypes, i.e. a higher Pareto’s beta (0.48)(Figure S13). Linkage disequilibrium values strongly departed from zero in both species, while *F*_IS_ was negative in *Cantharellus formosus* and positive in *Boletus edulis*, suggesting strong clonal rates (Stoeckel *et al*. 2021b). Overall, these low genotypic metrics (R, Pareto beta) and high genetic metrics (LD and *F*_IS_) allow us to characterize these populations as very clonal (c = 0.9 -1), giving an initial idea of clonal rates in natural populations of EM fungi.

Partial clonality and its impact on *N*_e_ estimation was explored by simulating clonality gradients (four conditions), specifically by increasing the number of replicated genotypes in five populations belonging to different species (hereafter: species)(Figure S10). Estimates of *N*_e_ with the LD method (Figure 2) were largely unreliable for the “H-W genotypes” (HW) condition, yielding infinite point estimates or confidence intervals in all species. In *Cantharellus formosus*, infinite point estimates or confidence intervals were observed in all four conditions. The SF method (Figure 2) yielded lower *N*_e_ point estimates than the LD method in all species and across all conditions, with *N*_e_ always lower than 100. Across all species, a pattern of decreasing *N*_e_ was observed as the level of clonality increased, especially in the populations with larger sample sizes, i.e. *Suillus luteus* and *Rhizopogon togasawarius*. *N*_e_ estimates across species were significantly lower in the LC condition than in all other conditions (Figure S14) when using the LD method and the SF method with the “random-mating” option. When using the “non-random mating” option, only the HW and LC conditions yielded significantly different values.

**Figure 2.**
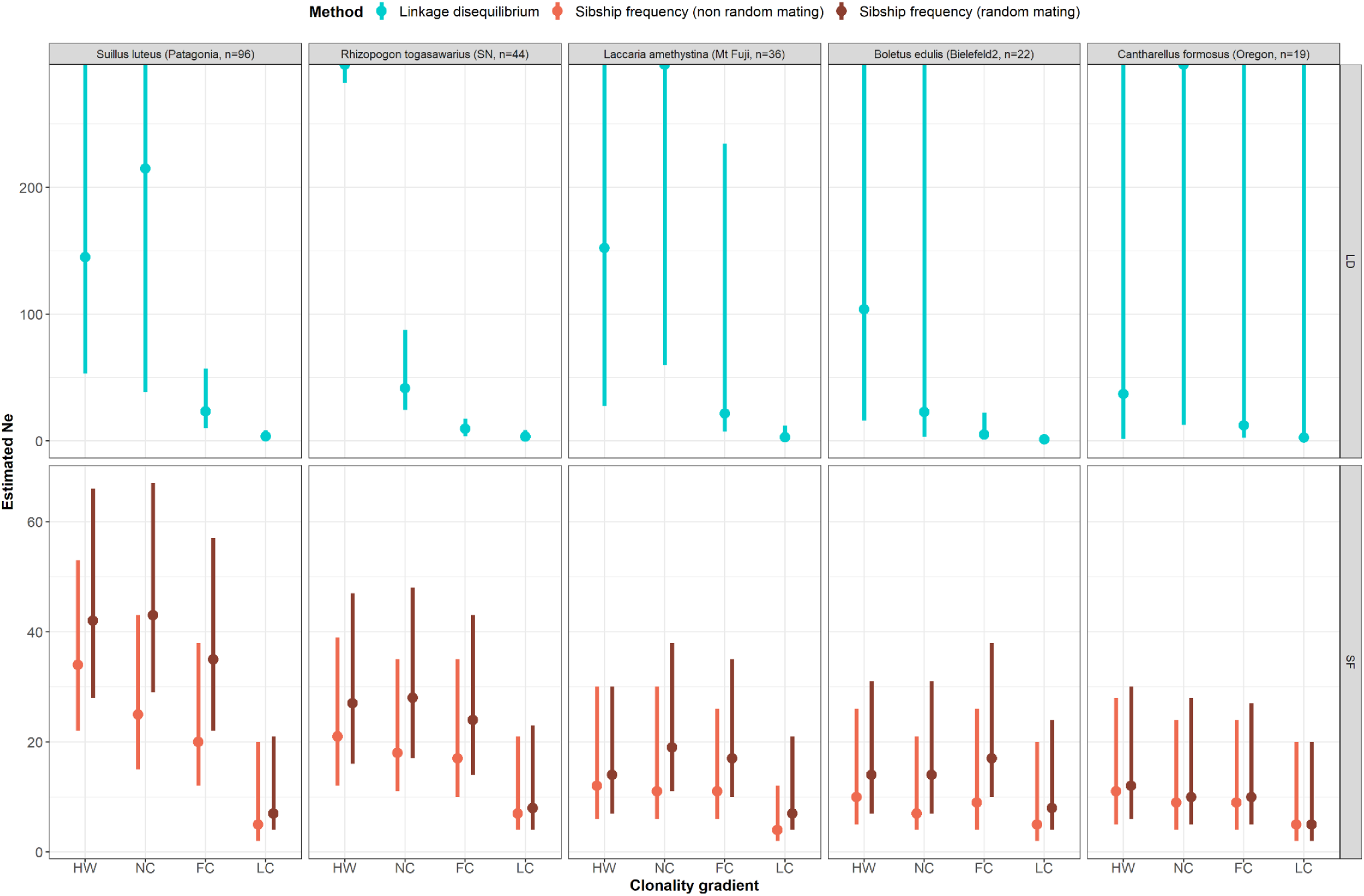
Estimation of *N*_e_ with the LD and SF methods in five populations belonging to different species of ectomycorrhizal fungi across an artificial clonality gradient consisting of four conditions: (1) genotypes are drawn randomly based on observed allelic frequencies (HW); (2) clone corrected data (NC); (3) original data with addition of 2-4 replicates per genotype (FC); (4) addition of one large clone to the original data (one genotype replicated 10-20 times) (LC). Each point represents the point estimate of *N*_e_ and the line range shows the Jackknife confidence interval (LD method) or the 95% confidence interval calculated from Student’s distribution (SF method). Points and lines extending beyond the plot represent cases where the *N*_e_ estimate or the upper limit are infinite.

### Influence of population structure on *N*_e_ estimations

We estimated *N*_e_ at increasing levels of genetic structure: at subpopulation level (sampling sites within a population), at population level (as defined in Table 2), and at regional and up to continental scale, by merging populations. *N*_e_ estimates and their confidence intervals are presented for *Rhizopogon togasawarius*, *Suillus brevipes* and *Laccaria amethystina* in Table 4. In most cases, LD and SF methods provided similar results, except for *Laccaria amethystina*, where negative results were obtained for almost all populations with the LD method.

**Table 4.**
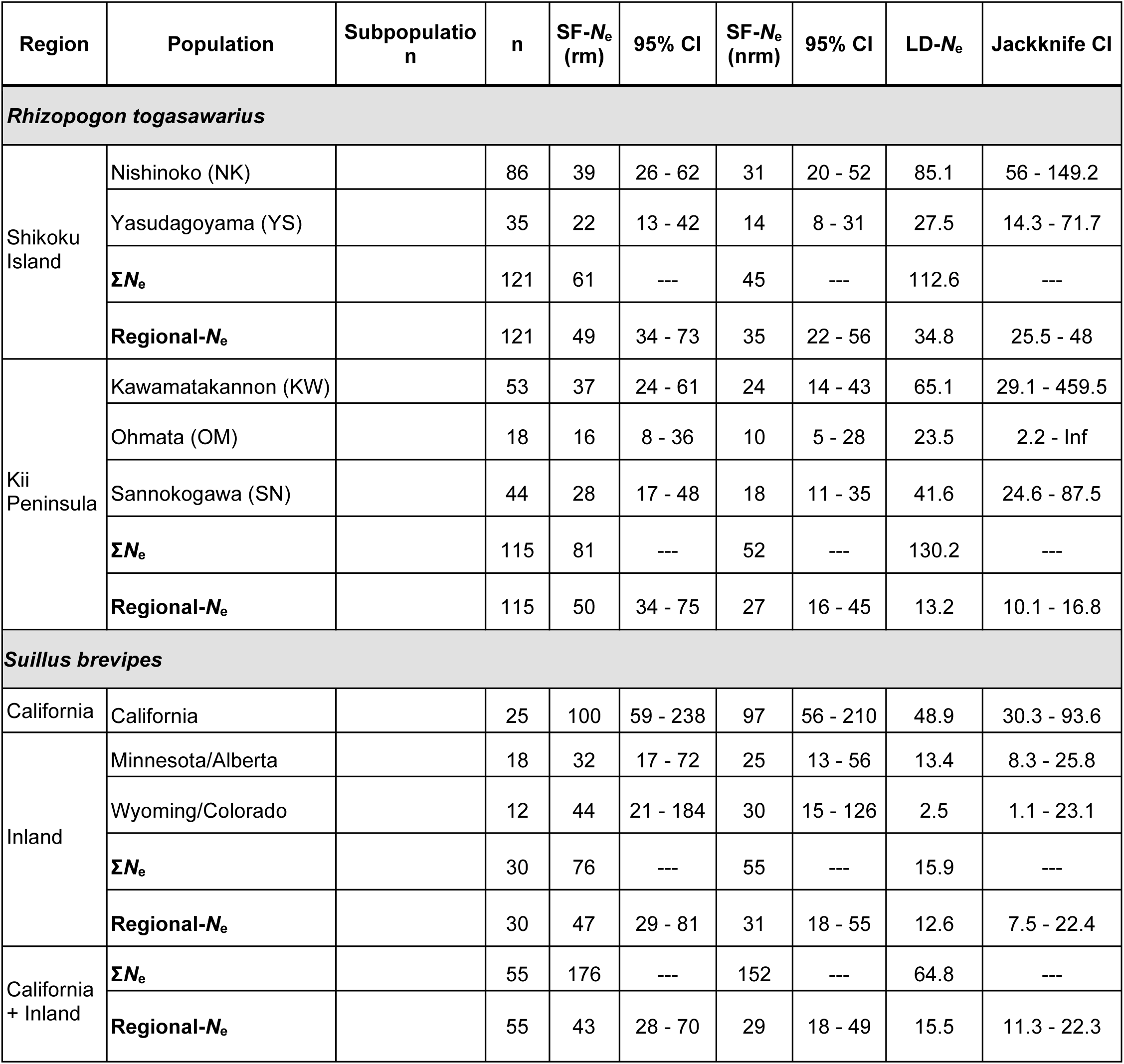
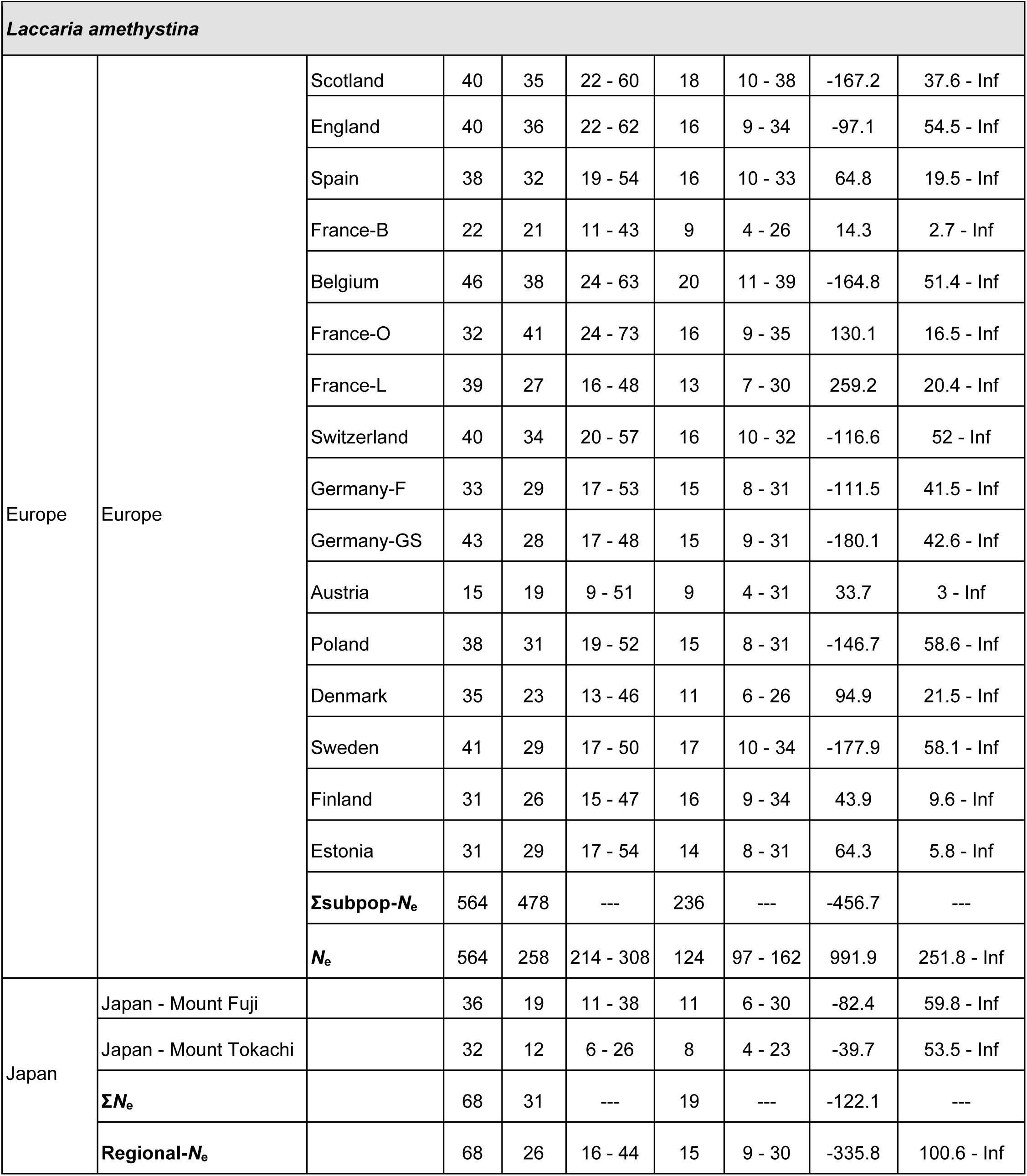
Estimation of *N*_e_ with the LD and SF methods at the regional (Regional-*N*_e_), population (*N*_e_) and subpopulation (subpop-*N*_e_) levels in *Rhizopogon togasawarius*, *Suillus bovinus* and *Laccaria amethystina*. In addition, sums of population level and subpopulation level estimates are provided for each region (∑*N*_e_ and ∑subpop-*N*_e_). Abbreviations: n, sample size; SF-Ne (rm), *N*_e_ estimation with SF method and “random mating” option; SF-*N*_e_ (nrm), *N*_e_ estimation with SF method and “non-random mating” option; LD-*N*_e_, Ne estimation with LD method; 95% CI, 95% confidence interval calculated from Student’s distribution; Jackknife CI, Jackknife confidence interval.

In *Rhizopogon togasawarius*, population-level *N*_e_ estimates ranged from 10 to 85.1 (Table 4), with systematically higher estimates (along with wider confidence intervals) with the LD method. Regional-level *N*_e_ estimates, conversely, were higher with the SF method. Regional-*N*_e_ estimates led to lower values compared to sums of population-level *N*_e_ estimates. This was true for both regions (Shikoku Island and Kii Peninsula), and for both methods, but the gap was much more pronounced for the LD method (values 3 to 10 times lower). In *Suillus brevipes,* local estimates ranged from 2.5 to 48.9 with the LD method, and from 25 to 100 with the SF method (Table 4). For both methods, higher estimates were observed in the California population (values between 48.9 and 100), while the other two populations had *N*_e_ estimates below 50. Regional-*N*_e_ estimate for Inland (Minnesota/Alberta + Wyoming/Colorado) was of similar magnitude compared to population-level *N*_e_ estimates. However, a larger grouping of populations at a continental level (Inland + California) resulted in lower estimates compared to the sum of population-level estimates (*N*_e_ = 15.5 - 43). In *Laccaria amethystina*, the population-level *N*_e_ estimate for Europe was of 258 with the SF method, which represented half of the total obtained by adding up all the subpopulations (Table 4). This effect was not very pronounced for the Japanese populations, where estimations from the regional level and sums of population-level *N*_e_ estimations were only marginally different.

### Influence of the number of markers used

The effect of the number of SNP markers on accuracy and precision of *N*_e_ estimates was assessed in the LD and SF methods using an increasingly larger number of SNPs from the three genomic datasets. Estimations of *N*_e_ are presented based on subsets of 2,000, 5,000, and 10,000 SNPs, with 10 replicates per population for *Suillus brevipes* (Figure 3), *Boletus edulis* (Figure S15), and *Suillus luteus* (Figure S16). Overall, an increase in the number of markers had no significant effect on the variance of the estimates (between replicates) and therefore accuracy was neither improved nor reduced. In a few populations (Europe-LD, East Coast-SF, Minnesota/Alberta-LD, Wyoming/Colorado - SF), *N*_e_ estimates were significantly more accurate when more markers were used, illustrated by the convergence of replicated point estimates along the gradient (Figure 3, Figure S15), although this effect was never observed for both methods when applied to the same population. Contrastingly, a better accuracy of the *N*_e_ estimates was observed when only 2,000 SNPs were used in the Europe population of *Boletus edulis*, with the SF method (Figure S15). The confidence intervals (CIs) of all *N*_e_ estimates did not vary across the markers gradient with both methods, except in two of the *Suillus brevipes* populations, where CIs become either wider (California) or narrower (Wyoming/Colorado) with an increase in the number of SNPs, only with the LD method (Figure 3). Overall, infinite CIs were observed in most cases, often alongside infinite or very high estimates (especially in *Suillus luteus*, Figure S16), suggesting that precision was not improved by an increase in the number of markers.

**Figure 3.**
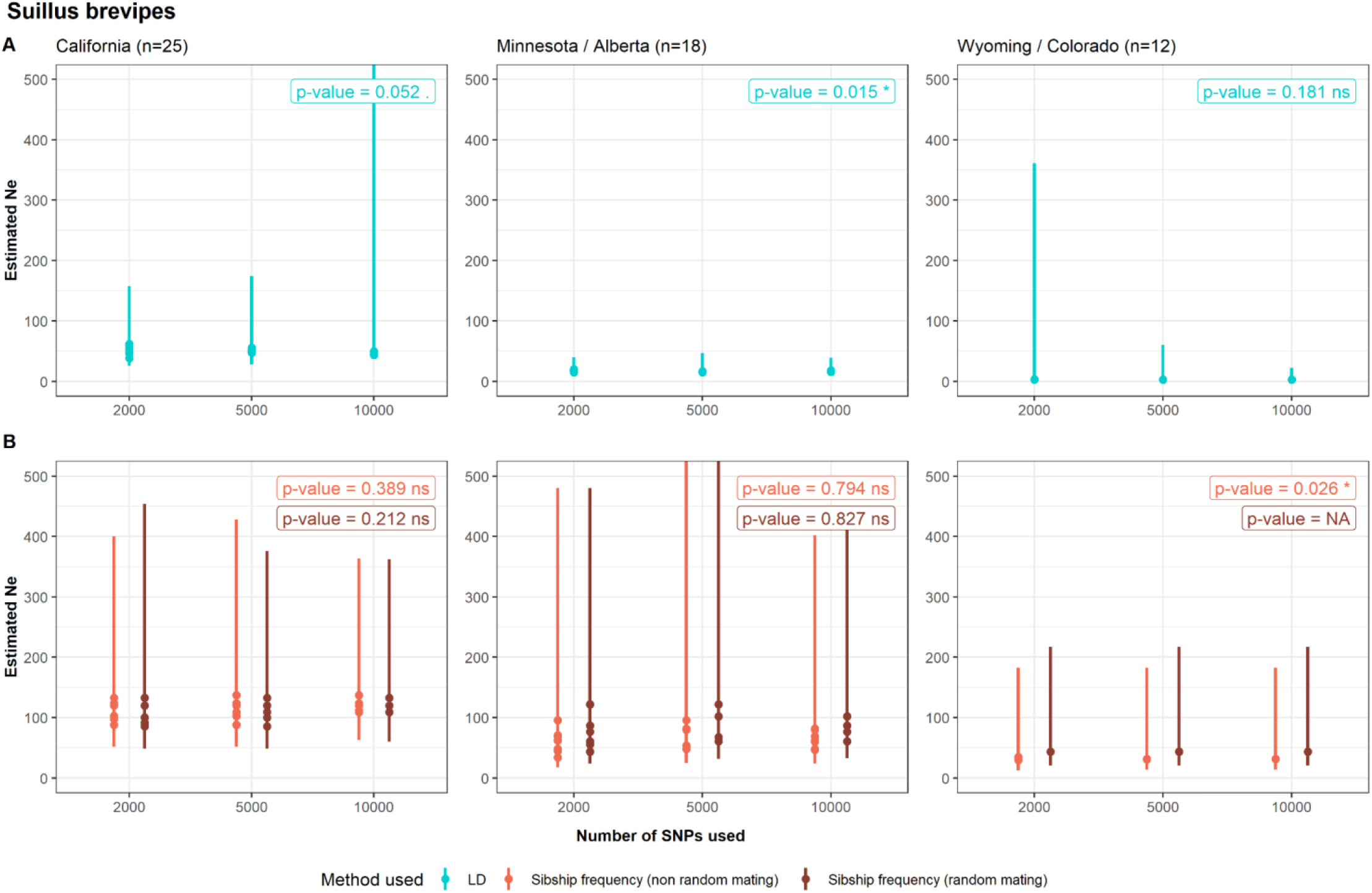
Estimations of *N*_e_ with LD method (A) and SF method (B) in 3 populations of *Suillus brevipes*. For each population, *N*_e_ is estimated based on 10 subsets of 2,000, 5,000 and 10,000 SNPs from the original datasets. Each point represents the value of *N*_e_ estimated by NeEstimator (LD) or Colony (SF), and the linerange shows the confidence interval (Jackknife confidence interval for LD method, 95% confidence interval calculated from Student’s distribution for the SF method). Points and lines that extend beyond the plot represent cases where the *N*_e_ estimate or the upper limit of the confidence interval is infinite. Note that the vertical axis is at different scales in panel B. P-values correspond to the results of a Levene’s test to assess the homogeneity of variances of the *N*_e_ estimates as a function of the number of SNPs used.

## Discussion

This study provides the first evaluation of effective population size estimation in ectomycorrhizal fungi. Leveraging genetic and genomic datasets extracted from the literature, we estimate the effective size of 19 populations across six species of ectomycorrhizal fungi. Using the LD-based method implemented in NeEstimator and the SF method implemented in Colony, we first provide an estimate based on population-level, clone-corrected datasets. We then evaluate how gradients of clonality, genetic structure and number of markers influence *N*_e_ estimates. Overall, both methods are affected by the presence of replicated genotypes, population structure, and variation in the number of markers used, but in directions and proportions that vary greatly from one dataset to another. Among the populations considered, only two of them have effective population sizes estimated to be above the *Ne500* threshold. However, these results should be taken with caution, and interpreted in the light of the various factors and violations of the *N*_e_ estimation assumptions examined in this study.

### Influence of partially clonal life cycles on *N*_e_ estimations

Using datasets containing a strong proportion of replicated genotypes, we assessed clonal rates based on genetic and genotypic metrics, following Stoeckel *et al*. (2021b). Our results suggest high clonality levels in *Cantharellus formosus* and, to a lesser extent, in *Boletus edulis* (Figures S12 and S13). This strong departure from random mating due to the presence of replicated genotypes runs counter to the assumptions underlying both estimation methods used here, with the caveat that small sample sizes and low marker resolution might reduce the reliability of the allelic frequencies and the clonal rates obtained (Arnaud-Haond *et al*. 2007).

To override these sampling limitations, we also studied *N*_e_ estimates along an artificial clonality gradient in five species. The clear decrease of the *N*_e_ estimates with increasing clonality gradient (Figure 2), particularly visible in the populations of *Suillus luteus* and *Rhizopogon togasawarius* and when using the LD method, can be explained by the introduction of additional LD in the datasets due to the increase in replicated genotypes. This reflects a “strong drift situation” leading to very low estimates of *N*_e_ (Stoeckel *et al*. 2021b; Waples 2024a). A similar, but less strong effect is observed with the SF methods in the same populations (Figure 2), which suggests that the SF method is not influenced by departures from random mating as much as the LD method is (Wang 2009). Here, the pattern can be explained by an increased relatedness due to the addition of replicated genotypes, with a lower *N*_e_ expected as the SF method is based on a measure of sibship. Overall, even if the two methods react to the increase in clonality with different magnitudes, significantly lower *N*_e_ estimates are consistently observed when a very large clone is simulated in the datasets (Figure S14): the introduction of a frequently occurring genotype leads to a dramatic increase in the level of LD or of relatedness, as a large proportion of individuals become similar in the dataset.

Nevertheless, the pattern of decreasing *N*_e_ estimates with increasing clonality is not visible in all EM fungal species, especially when using the SF method. In addition, *N*_e_ estimates obtained from the SF method are consistently lower than 100, whereas the LD estimation yields many infinite values. The lower values obtained from the SF method can be explained by a tendency of this method to underestimate *N*_e_ when (i) the actual *N*_e_ is large, (ii) sample size is small and (iii) only a few markers are used (Wang 2009). This might be the case for some of our microsatellite datasets (number of markers 5-20, sample sizes 19-96), even though the actual *N*_e_ of these populations remains unknown. Similarly, a lack of genetic drift signal is known to cause infinite estimates when using the LD method, for instance when trying to estimate a large *N*_e_ using a small set of samples (Waples 2024a) or when marker resolution is lacking. This is the case with the *Cantharellus formosus* dataset, which contains 19 different genotypes obtained from only 5 SSR markers, and where all estimates of *N*_e_ across the clonality gradient are infinite (Figure 2). Interestingly, clearer patterns of decreasing *N*_e_ across the clonality gradient are observed in populations with larger sample sizes. All this put together proves the limits of small datasets when using both estimation methods, especially if the expected *N*_e_ is large, as previously reported (Wang 2025; Waples 2021).

The datasets considered in this study may not be directly informative of the effect of partial clonality on *N*_e_, for instance they do not allow us to test whether partially clonal populations have smaller effective population sizes than purely sexual populations. This latter issue might require a more theoretical approach or several empirical datasets collected ad hoc. However, our results show that the presence of replicated genotypes introduces a downward bias in the estimation of *N*_e_, with stronger effects when using the LD method.

### Accounting for population structure in *N*_e_ estimations

The effect of population genetic structure on *N*_e_ estimation was assessed by analysing the datasets containing very differentiated populations or wide-range samplings of *Rhizopogon togasawarius, Suillus brevipes* and *Laccaria amethystina*. In most cases, the admixture of two or more genetically distinct populations led to a downwardly biased *N*_e_ estimate, especially with the LD method. This bias has already been described in previous studies, and results from mixture LD due to a two-locus Wahlund effect (Mergeay *et al*. 2024; Waples 2024a; Waples & England 2011). In other words, pooling offspring from genetically differentiated groups of parents generates non-random allele co-occurrence patterns across individuals, some combinations of alleles will never be observed simultaneously in the same individual, while others are over-represented. The LD-method will interpret this linkage as a signal of genetic drift, and underestimate *N*_e_ (Neel *et al*. 2013; Waples & England 2011).

In *Rhizopogon togasawarius* (Table 4), where 5 strongly differentiated populations have been described (Abe *et al*. 2024), *N*_e_ estimates from the LD method (and SF method to a lesser extent) were systematically lower when different populations are combined together prior to the estimation, compared to *N*_e_ obtained from the sum of population-level estimates. Thus, estimating *N*_e_ from pooled populations does not reflect the sum of the local effective population sizes (Fedorca *et al*. 2024; Waples 2024a), because of the mixture LD bias. While this bias is evident with the LD method, the SF method produced similar estimates or upwardly biased values when several populations were pooled, potentially caused by a decrease in relatedness among individuals when very divergent populations are mixed. Results for *Suillus brevipes* were more nuanced (Table 4) with a downward bias only observed with the LD method at the continental level (when very differentiated populations are grouped), while the regional estimates (combination of two populations) corresponded to the average over the population level estimates. The SF method, on the other hand, seems unaffected by the different groupings, as values remain consistent across scales. The relative difference in dispersal ability and thus in population isolation between *S. brevipes* and *R. togasawarius* might be an explanation of why genetic structure has a lesser impact on *N*_e_ estimation in *S. brevipes*. In *S. brevipes* the downward bias is not observed at a finer scale probably because its dispersal capacity through aerial spores is significant; *R. togasawarius*, on the other hand, produces hypogeous and animal-dispersed spores, making fine-scale genetic structure more discrete.

In the case of *Laccaria amethystina*, only SF-based estimations can be properly interpreted, as results from the LD method led to negative or infinite values (Table 4). Such values might be indicative of very large population sizes, which is consistent with previous studies in *Laccaria amethystina*. According to Vincenot *et al*. (2012), a single panmictic population is occurring across Europe, with no isolation by distance (IBD) across > 2900km. Moreover, high diversity of small genets has been observed at the local scale (Fiore-Donno & Martin 2001), and the repeated settlement of new spores every year. These two aspects are likely to strongly limit genetic drift, and as a consequence, high *N*_e_ are expected in this species (Gherbi *et al*. 1999). Surprisingly, *N*_e_ estimations from the SF method were much lower than we might expect (population level estimates < 50) and showed a clear downward bias when all subpopulations of Europe were combined prior to the estimation, compared with the estimation obtained from the sum of each subpopulation’s *N*_e_ estimate (Table 4). This suggests the presence of some discrete genetic clusters in the European cluster, or at least a weak genetic structure associated with recent gene flow that might bias *N*_e_ estimation using the SF method.

Using three contrasting species’ datasets, we can outline answers to both methodological and biological questions concerning the impact of genetic structure on *N*_e_ estimation. First, our results show significant and expected biases on the *N*_e_ estimates when they are computed on incorrectly delimited populations, for instance the downward bias in LD-*N*_e_ estimates resulting from mixture LD. Concerning the SF method, we demonstrate that some ectomycorrhizal fungal species with long-distance dispersal might display high gene flow that could bias *N*_e_ estimation, although to a lesser extent than with the LD method. In addition, the presence of negative or infinite values suggesting very large populations (such as in *Laccaria amethystina*) places serious constraints on the use of these tools to properly estimate *N*_e_ when having limited sampling and marker resolution. Among the new methods that are being developed to account for genetic structure when estimating *N*_e_, the one implemented in CurrentNe2 enables more accurate estimates in large and continuously distributed populations (Santiago *et al*. 2025) and has already been applied in tree populations (Caballero *et al*. 2026). This method seems promising for *N*_e_ estimation in ectomycorrhizal fungal populations, which are known to display wide distribution ranges and patterns of isolation by distance. This method being only available for SNPs datasets mapped to a reference genome, it could unfortunately not be applied to datasets based on microsatellite genotyping.

### Use of genomic datasets for *N*_e_ estimation

As pseudoreplication issues are expected when using genomic-scale datasets (Waples 2024a), we tested whether the number of markers used would affect the accuracy and the precision of *N*_e_ estimates. We observe no clear pattern of increasing or decreasing accuracy of the estimates when the number of markers is increased. However, a few exceptions appear in both methods, with accuracy increasing when a larger number of markers is used, but never simultaneously for both methods within the same population (Figure 3). In terms of precision, many simulation works have highlighted a potentially reduced precision in the *N*_e_ estimates, pictured by large confidence intervals when using genomic-scale datasets (Waples 2021; Waples *et al*. 2022). Here, contrary to expectations, no significant effect of the number of markers on the precision of *N*_e_ estimates was observed, neither for the LD nor the SF method. As a caveat, the prevalence of infinite values and confidence intervals might suggest that some populations are very large and therefore impossible to measure with a limited sampling size and the analytical tools available, for instance in *Suillus luteus* (Figure S16), where only 34 individuals were sampled.

### Recommendations and perspectives for conservation genetics

Although *N*_e_ is widely recognized as a key evolutionary and conservation metric in most taxa (Hoban *et al*. 2021b), we show that existing estimators such as those based on the LD and SF methods are associated with several challenges arising from the biological characteristics of EM fungi, as well as methodological aspects of genetic and genomic datasets. Thus, until the biological factors influencing *N*_e_ and the sources of methodological bias will be clarified, the *Ne500* indicator cannot be considered directly applicable to EM fungi. Some of the drawbacks illustrated here are common to other species with large continuously distributed populations, and advances in genomic methods for *N*_e_ assessment are already emerging, in trees for example (Caballero *et al*. 2026). Two other factors affecting the estimation of *N*_e_ could not be tested here but remain of importance. First, both the LD and the SF method assume non-overlapping generations, i.e. the study of a single cohort, a condition that is very unlikely to be met, given that ectomycorrhizal fungi can be perennial in the soil and produce fruitbodies over several years. Secondly, this study uses datasets extracted from the literature that rely both on microsatellite and SNPs markers. Our results show that most summary statistics are homogeneous across populations from the same datasets, but that there is some discrepancy when comparing populations from the same species analysed with different markers (SNPs vs SSRs). Thus, we need to remain cautious when comparing datasets with different types and number of markers.

Other *N*_e_ estimation issues appear specific to the fungal species analysed here, and would necessitate more investigation to gain a good understanding of what shapes genetic diversity in EM basidiomycete fungi. The evolutionary consequences of dispersal patterns, rates of clonality, census sizes, and the relative importance of haploid and dikaryotic phases are some of the factors that could be explored both independently, but also in combination in eco-evolutionary simulation scenarios. As an example, we already identified that the genetic structure of populations is closely linked to the dispersal mode of each species (wind or animal dispersed spores), along with the balance between sexual and asexual reproduction. On the other hand, the design of population genetics studies needs to be redefined in order to better encompass the genetic diversity of the whole population. Most studies rely on genetic data from only one compartment, fruitbodies or mycelium. However, genetic individuals can produce several fruitbodies in the same season, or no fruitbody at all during their life, making this type of sampling certainly incomplete (Vincenot & Selosse 2017). A joint sampling of fruitbodies and soil or root-tip mycelium would be the ideal strategy to obtain a more accurate picture of the current genetic diversity in the whole population, and could thus be compared with the simulated expectations.

Ectomycorrhizal fungal species are key components of most forest ecosystems, and their integration in conservation genetic programs is still limited compared to other taxa, because of the challenges associated with their biology explored above. Some of the species considered in this study are classified as “endangered” by the IUCN Red List, such as *Rhizopogon togasawarius* (Murata & Nara 2022), with a few thousands individuals remaining, while others have a “least concern” status with varying levels of knowledge on their current population trends (Dahlberg 2019a, b; Mueller 2025; Siegel 2021), or are even not evaluated (e.g. *Suillus brevipes*). Our study represents the first step towards assessing the feasibility of applying the existing genetic indicator *Ne500* in support of current conservation guidelines and efforts. We expect that the methodological avenues we suggested above will help improve our understanding of population genetic diversity in ectomycorrhizal fungi and verify the applicability of existing and new genetic indicators for fungal conservation.

## Acknowledgements

This study was supported by government funding administered by the French National Research Agency (ANR) under the France 2030 initiative, reference number ‘ANR-24-RRII-0003’, and implemented through the INRAE EXPLOR’AE programme. We would like to thank the authors who shared their genetic and genomic data with us, in particular Susie Dunham (Oregon State University, USA), Keaton Tremble (Duke University and University of Illinois, USA) and María Belén Pildain (National Scientific and Technical Research Council, Argentina). We are grateful to the genotoul bioinformatics platform Toulouse Occitanie (Bioinfo Genotoul, https://doi.org/10.15454/1.5572369328961167E12) for providing computing and storage resources. We would like to thank Marie-Gabrielle Harribey (INRAE, France) for the help provided in the implementation of NeEstimator in R, and Olivier Lepais (INRAE, France) and Irene Martinez Velasco (National Autonomous University of Mexico, Mexico) for helpful contributions to the development of an automated workflow for Colony.

## Author contribution statement

MH, RG, AC designed the study. MH, RG supervised the project. AC, RG carried out the literature search for datasets. AB provided genomic data for the analyses. AC processed and analysed the data. AC, MH and RG wrote the manuscript. All authors contributed with edits and suggestions for the analyses and manuscript.

## Data archiving

Original datasets are available at: https://doi.org/10.5281/zenodo.21072336 Detailed scripts will be available upon publication.

