## Supplementary Figures for "Towards genetic indicators in ectomycorrhizal fungi: estimating the effective population size"

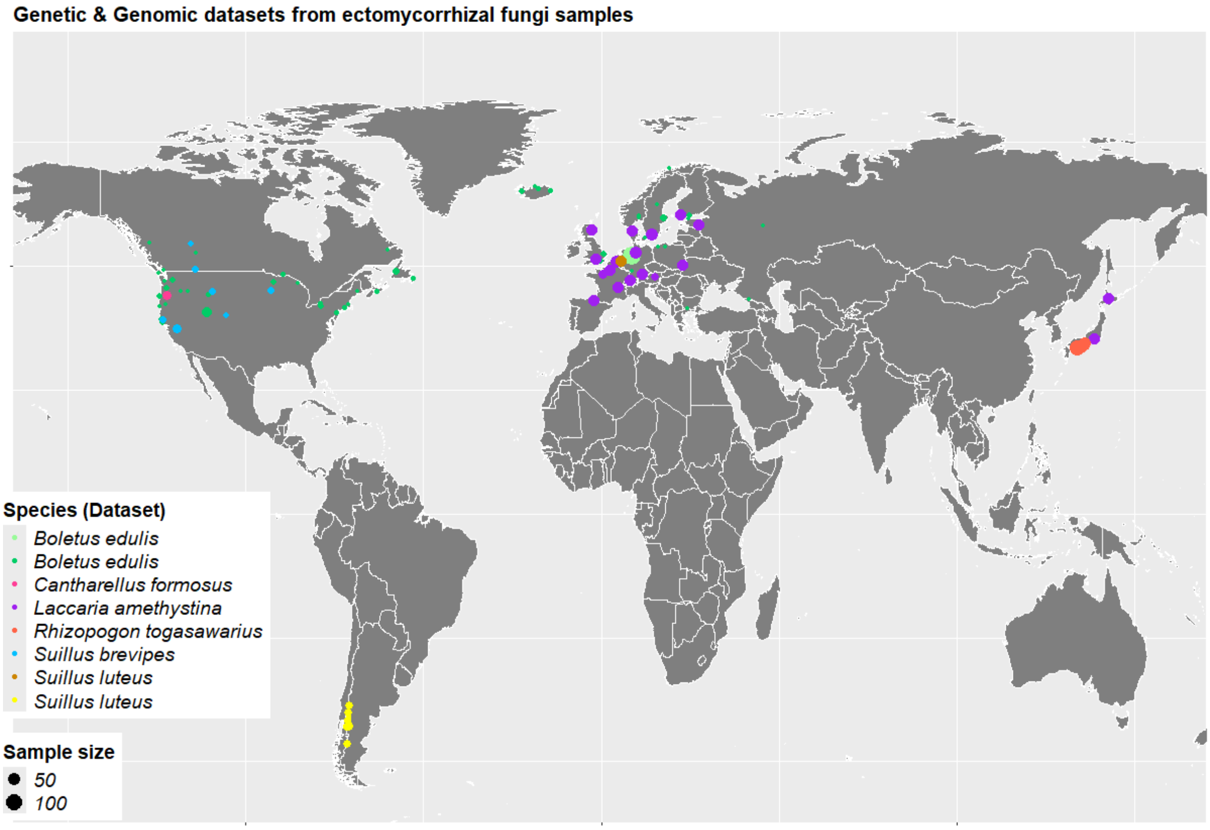

Figure S1. Map of the sampling points from the 8 datasets described in Table 2.

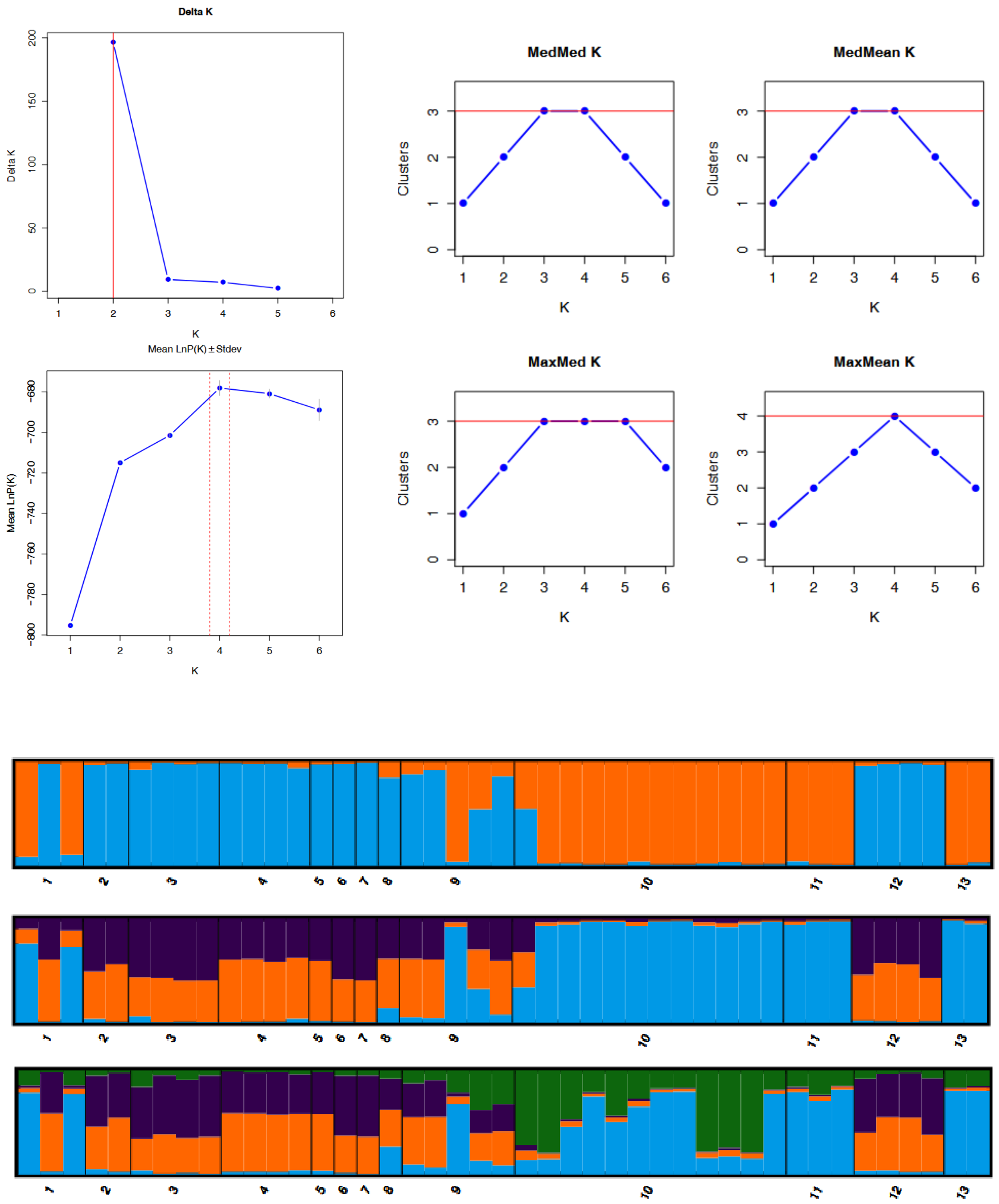

Figure S2. Results of STRUCTURE analysis for the *Boletus edulis* (Hoffman et al. 2020) dataset. Outputs from StructureSelector: K estimators (Delta K, LnP(K), MedMed K, MedMean K, MaxMed K, MaxMean K) and barplots (K = 2 to K = 4).

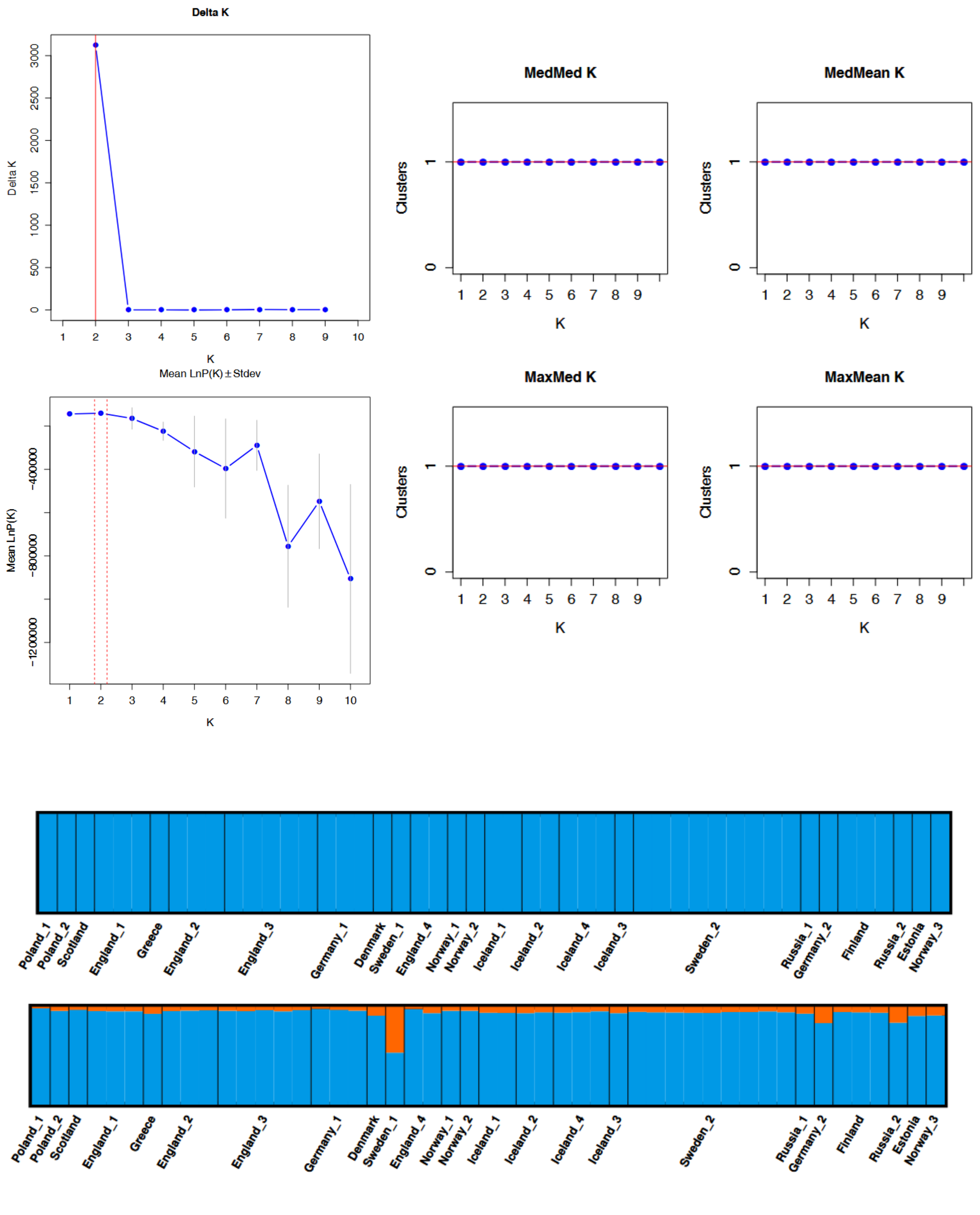

Figure S3. Results of STRUCTURE analysis for the *Boletus edulis* (Brejon Lamartinière *et al.* 2024) dataset - Europe Lineage (EU). Outputs from StructureSelector: K estimators (Delta K, LnP(K), MedMed K, MedMean K, MaxMed K, MaxMean K) and barplots (K = 1 to K = 2).

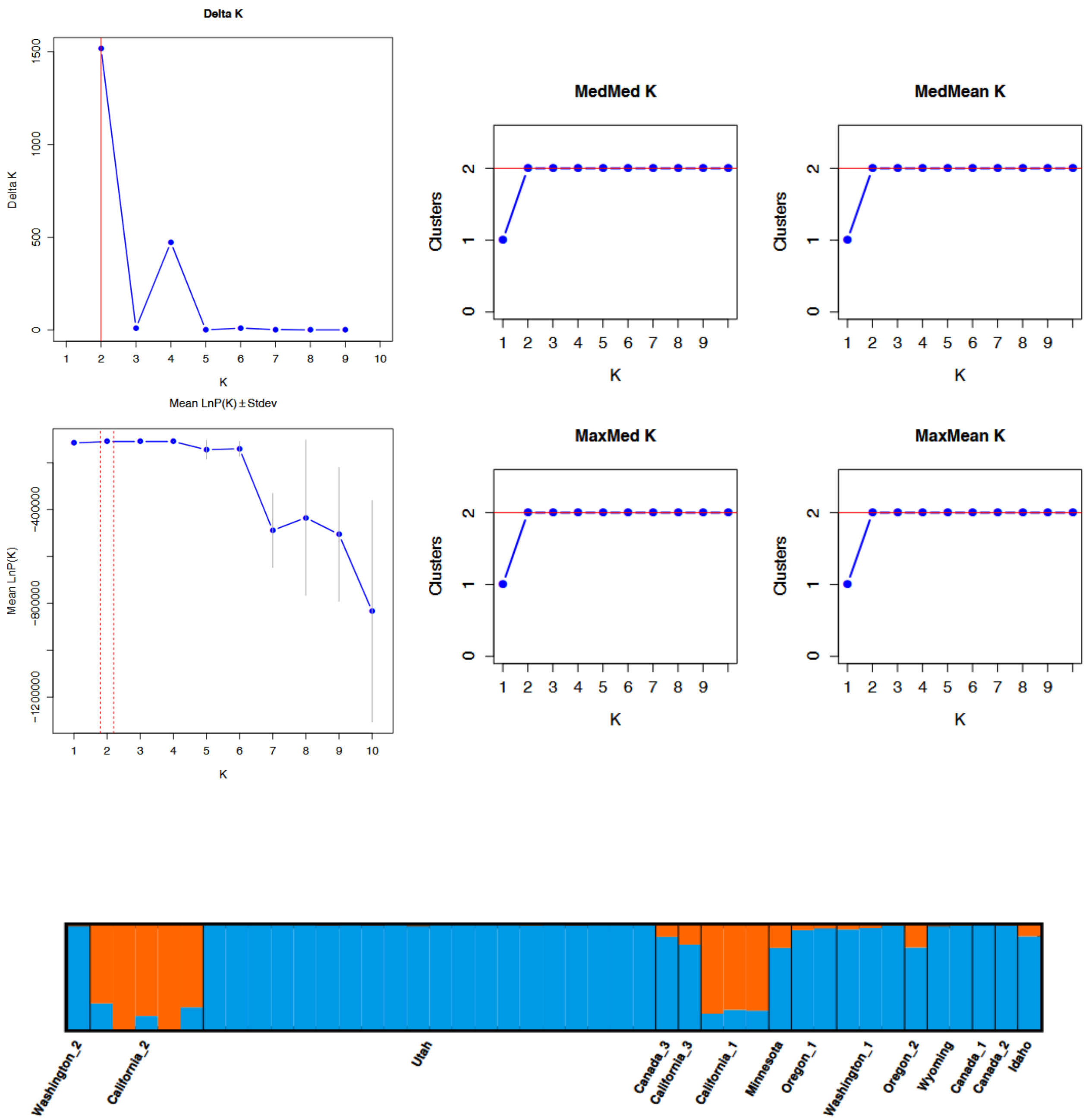

Figure S4. Results of STRUCTURE analysis for the *Boletus edulis* (Brejon Lamartinière *et al.* 2024) dataset - West Coast Lineage (WC). Outputs from StructureSelector: K estimators (Delta K, LnP(K), MedMed K, MedMean K, MaxMed K, MaxMean K) and barplots (K = 2).

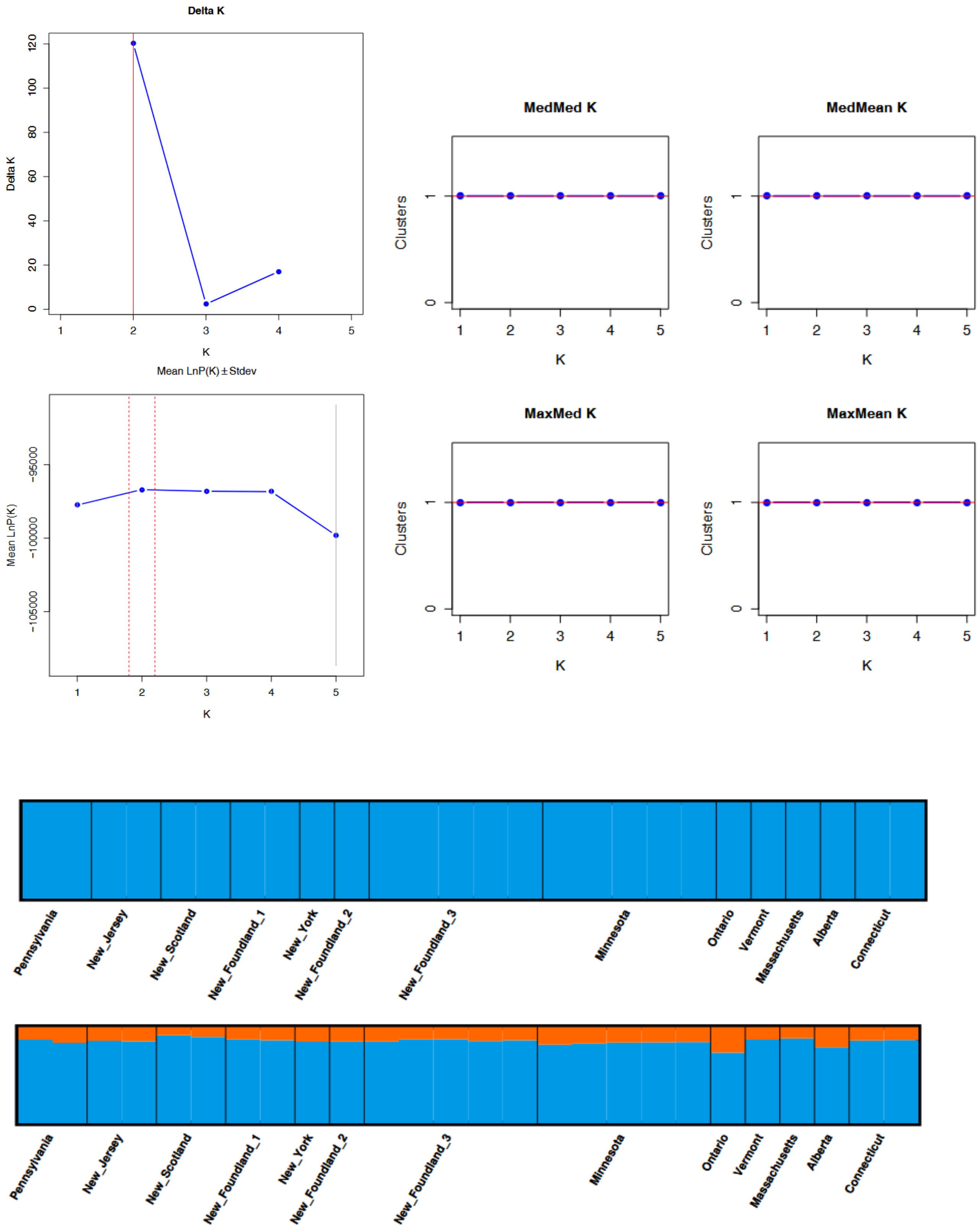

Figure S5. Results of STRUCTURE analysis for the *Boletus edulis* (Brejon Lamartinière *et al.* 2024) dataset - East Coast Lineage (EC). Outputs from StructureSelector: K estimators (Delta K, LnP(K), MedMed K, MedMean K, MaxMed K, MaxMean K) and barplots (K = 1 to K = 2).

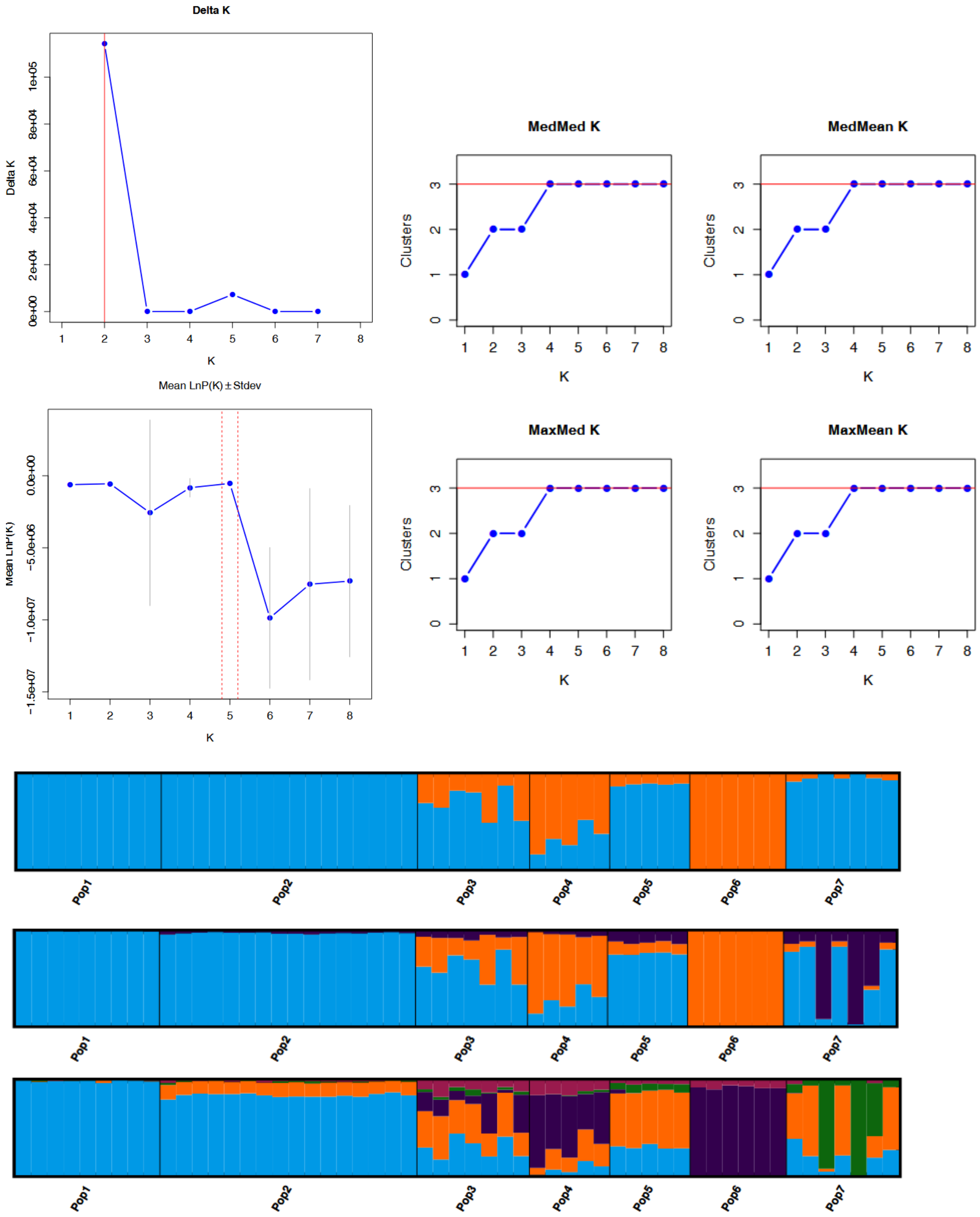

Figure S6. Results of STRUCTURE analysis for the *Suillus brevipes* (Branco et al. 2017) dataset. Outputs from StructureSelector: K estimators (Delta K, LnP(K), MedMed K, MedMean K, MaxMed K, MaxMean K) and barplots (K = 2, K = 3, K = 5).

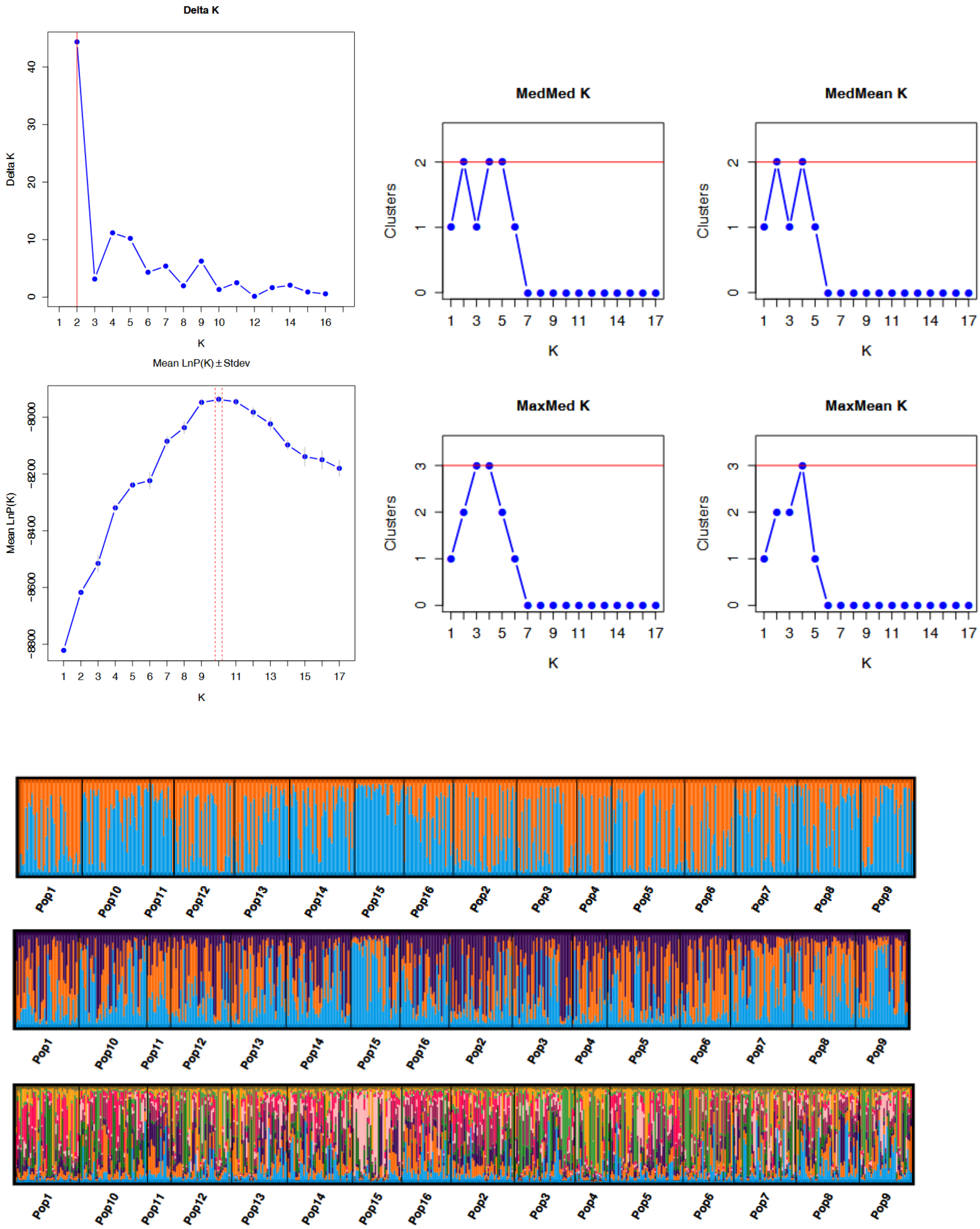

Figure S7. Results of STRUCTURE analysis for the *Laccaria amethystina* (Vincenot et al. 2012) dataset - Europe cluster. Outputs from StructureSelector: K estimators (Delta K, LnP(K), MedMed K, MedMean K, MaxMed K, MaxMean K) and barplots (K = 2, K = 3, K = 10).

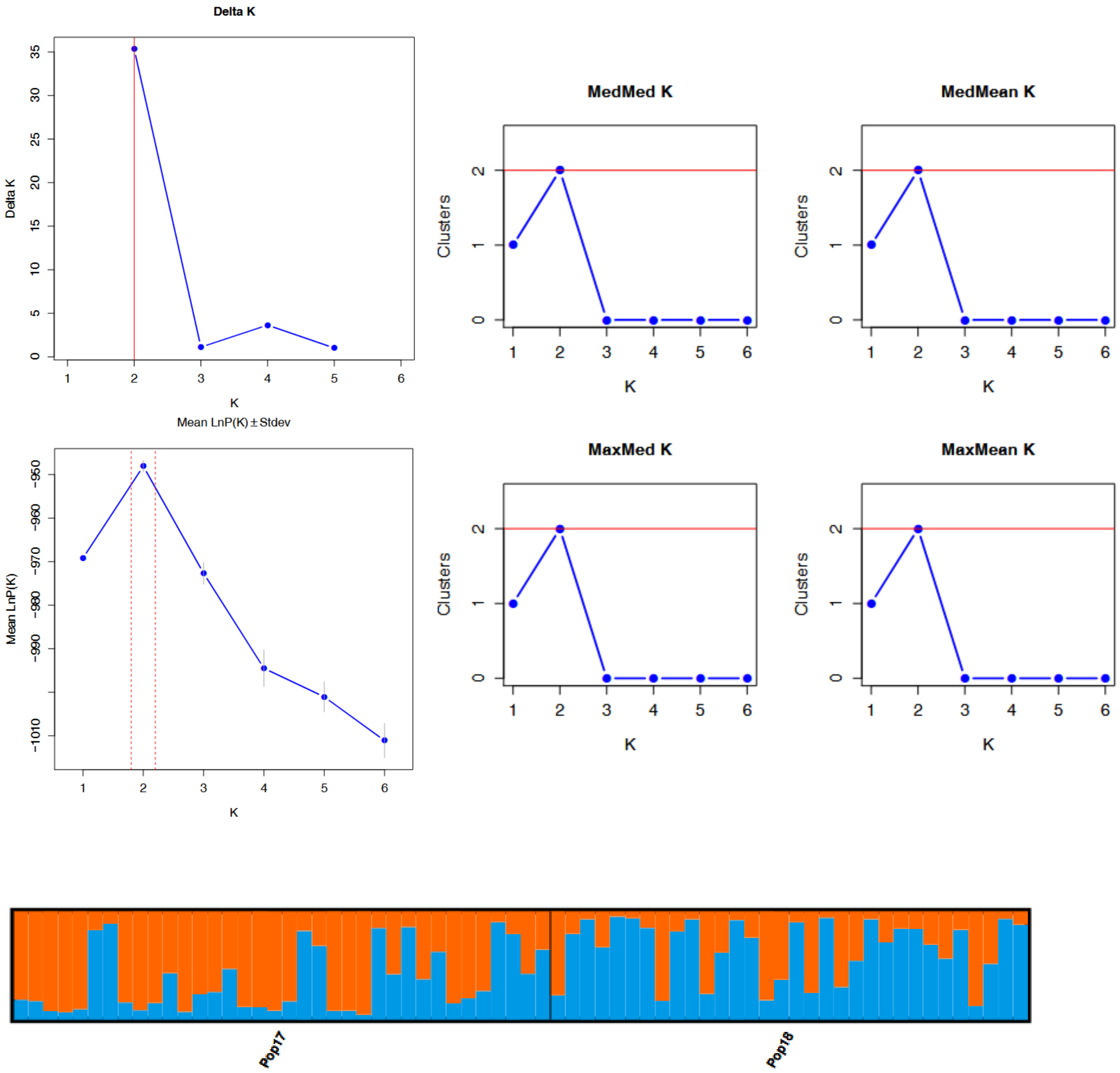

Figure S8. Results of STRUCTURE analysis for the *Laccaria amethystina* (Vincenot et al. 2012) dataset - Japan cluster. Outputs from StructureSelector: K estimators (Delta K, LnP(K), MedMed K, MedMean K, MaxMed K, MaxMean K) and barplots (K = 2).

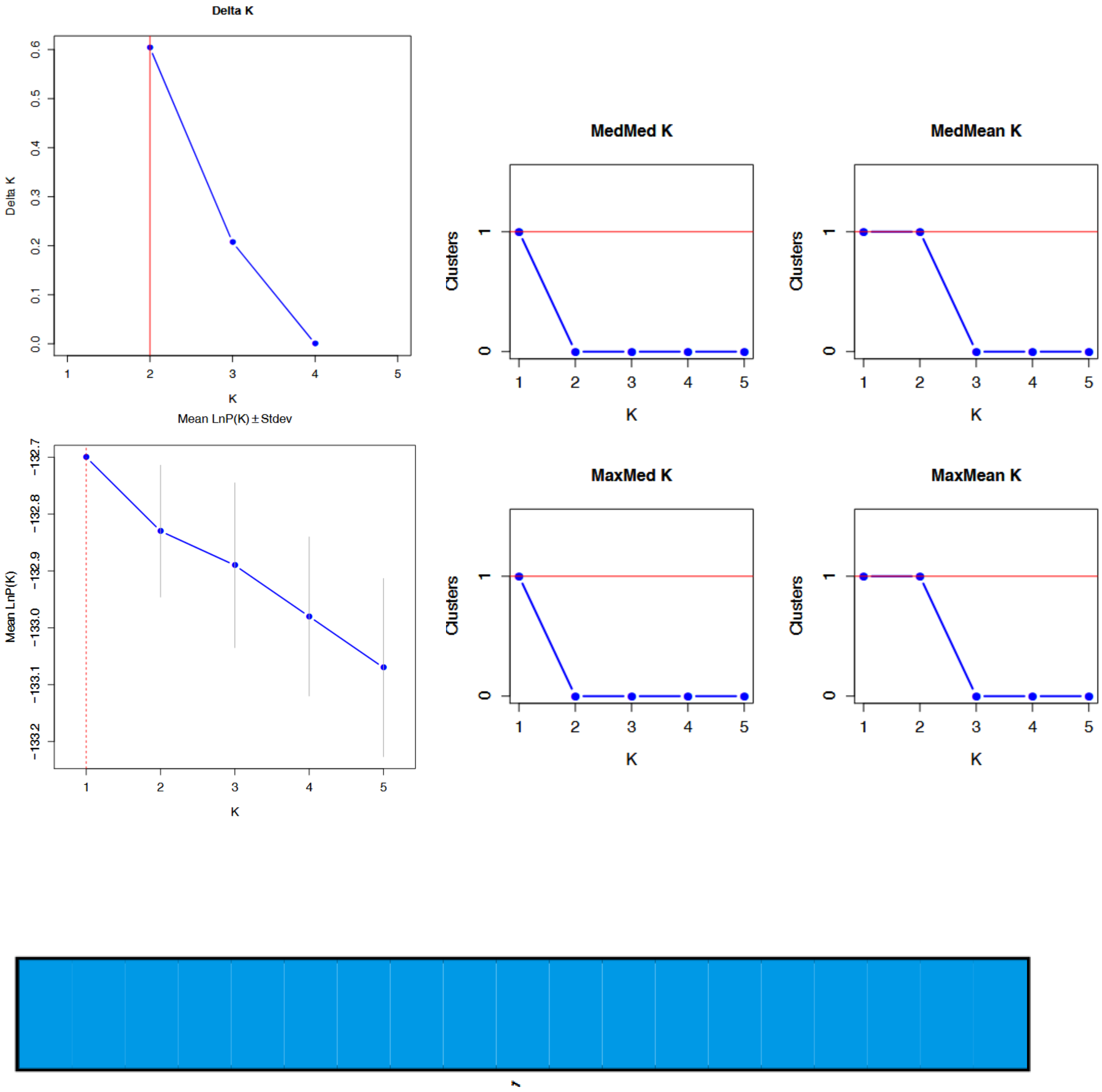

Figure S9. Results of STRUCTURE analysis for the *Cantharellus formosus* (Dunham et al. 2006) dataset. Outputs from StructureSelector: K estimators (Delta K, LnP(K), MedMed K, MedMean K, MaxMed K, MaxMean K) and barplots (K = 1).

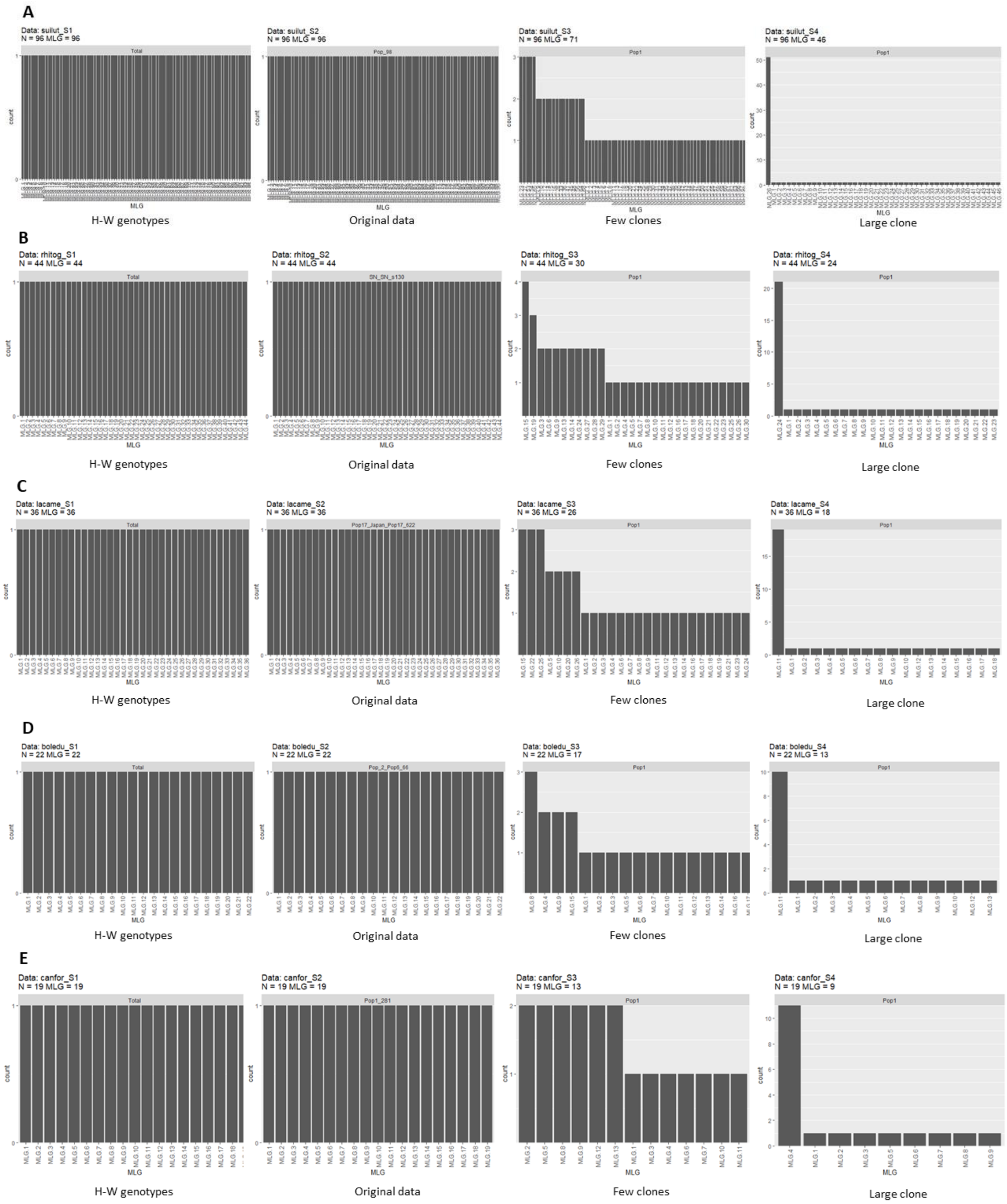

Figure S10. MLG plots for each of the 4 datasets composing the clonality gradient in 5 populations of ectomycorrhizal fungi : *Suillus luteus* - Pop Patagonia (A), *Rhizopogon togasawarius* - Pop Sannokogawa (B), *Laccaria amethystina* - Pop Mount Fuji (C), *Boletus edulis* - Pop Bielefeld_2 (D), *Cantharellus formosus* - Pop Oregon (E).

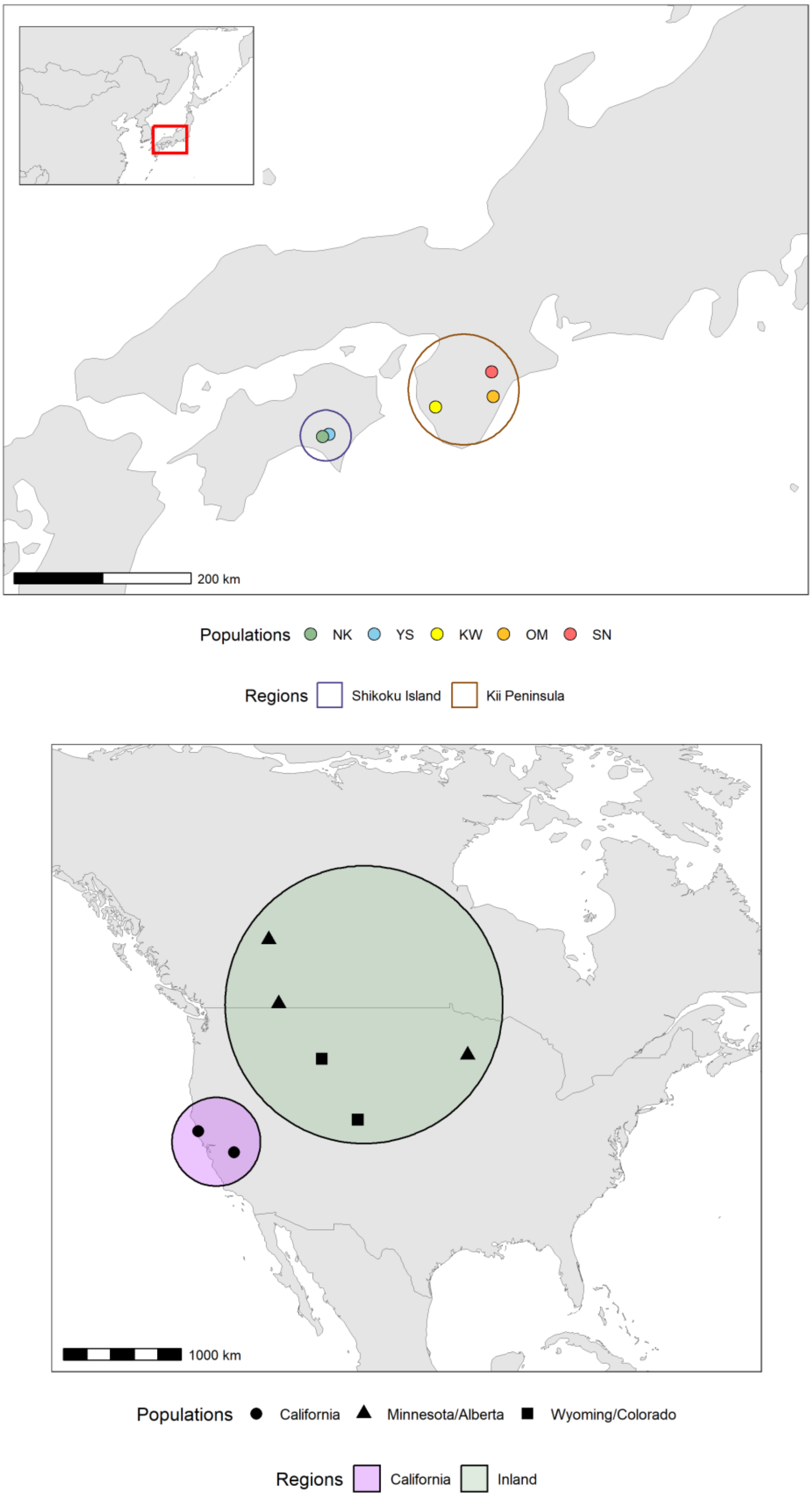

Figure S11. **(Top)** Map of the sampling points and corresponding populations and regions for *Rhizopogon togasawarius* (adapted from Abe *et al.* 2024). **(Bottom)** Map of the sampling points and corresponding populations and regions for *Suillus brevipes* (adapted from Branco *et al.* 2017).

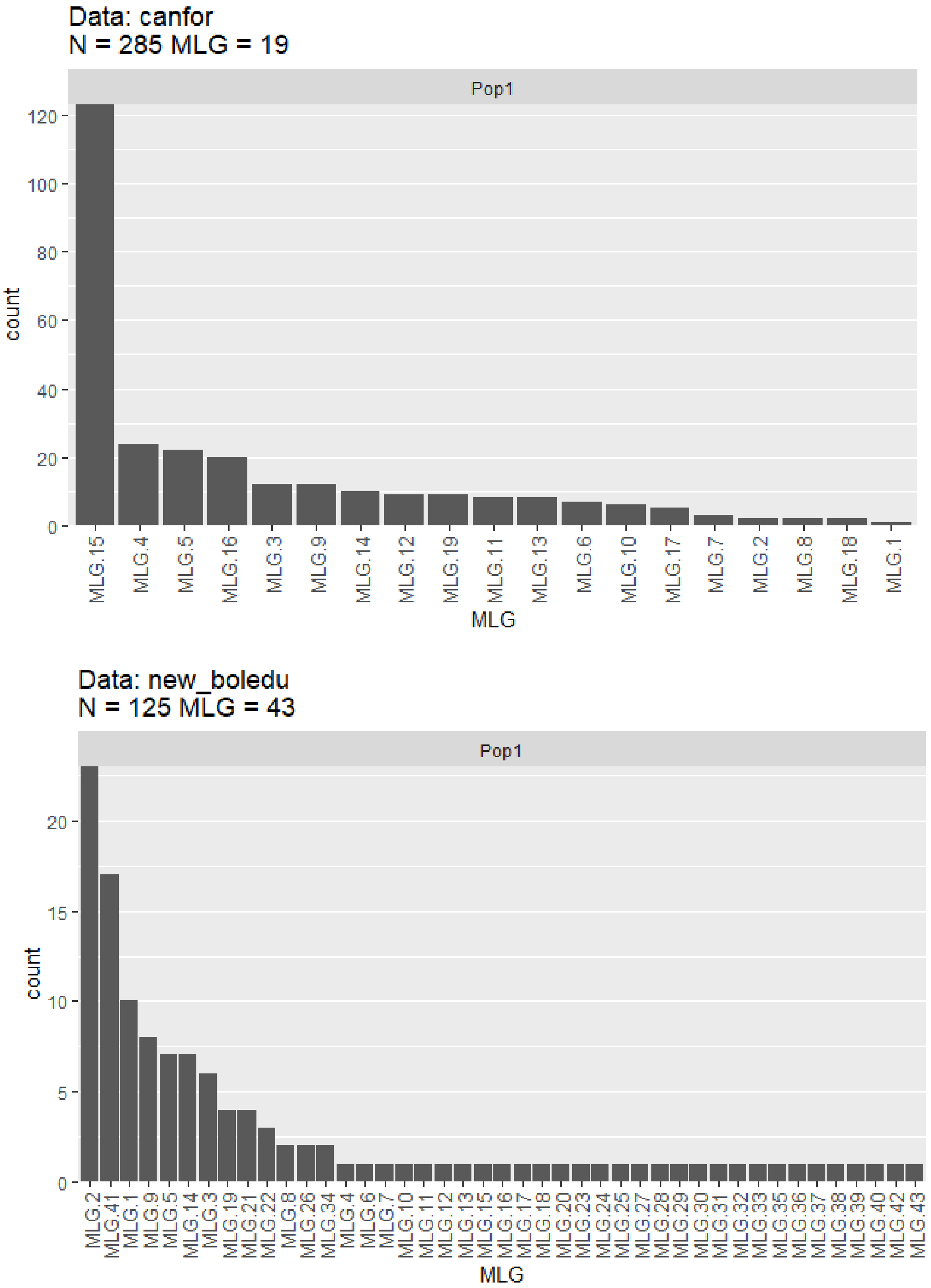

Figure S12. MLG plots for two datasets: *Cantharellus formosus* (top) and *Boletus edulis* (bottom). The datasets used for this analysis are not clone corrected, and assume all samples are part of the same population.

| | R | Pareto's beta | $F_{IS}$ | rd |
|---|---|---|---|---|
| ***Cantharellus formosus*** | 0.06 | 0.2 | -0.09 | 0.247 ** |
| ***Boletus edulis*** | 0.34 | 0.48 | 0.19 | 0.214 ** |

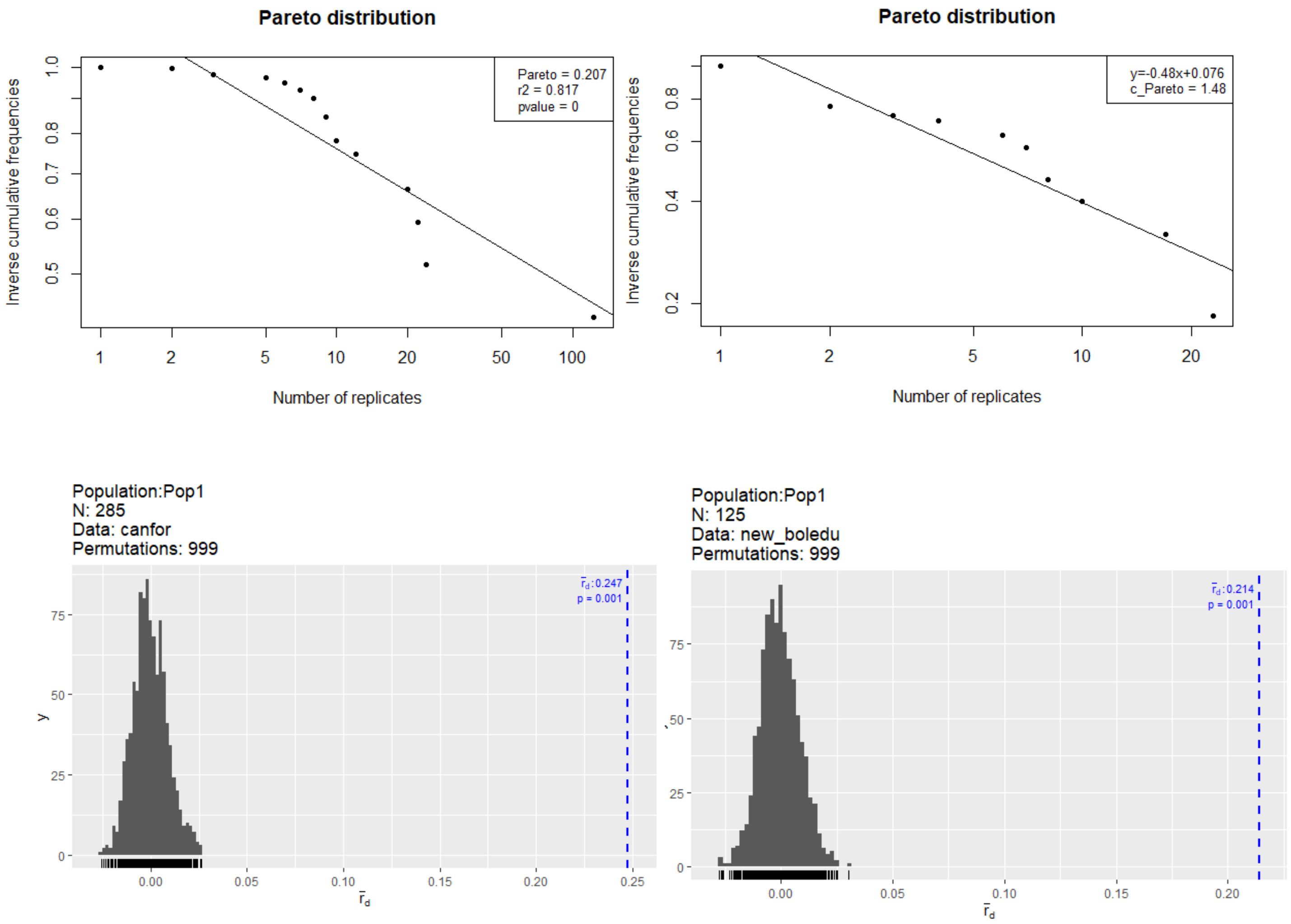

Figure S13. Genetic and genotypic indices for two datasets : *Cantharellus formosus* and *Boletus edulis*. Abbreviations: R, Genotypic richness; $F_{IS}$, inbreeding coefficient; rd, Linkage disequilibrium; ** indicate significant values ($p < 0.01$). Pareto's beta and rd plots for both datasets are shown below (*Cantharellus formosus* on the left and *Boletus edulis* on the right).

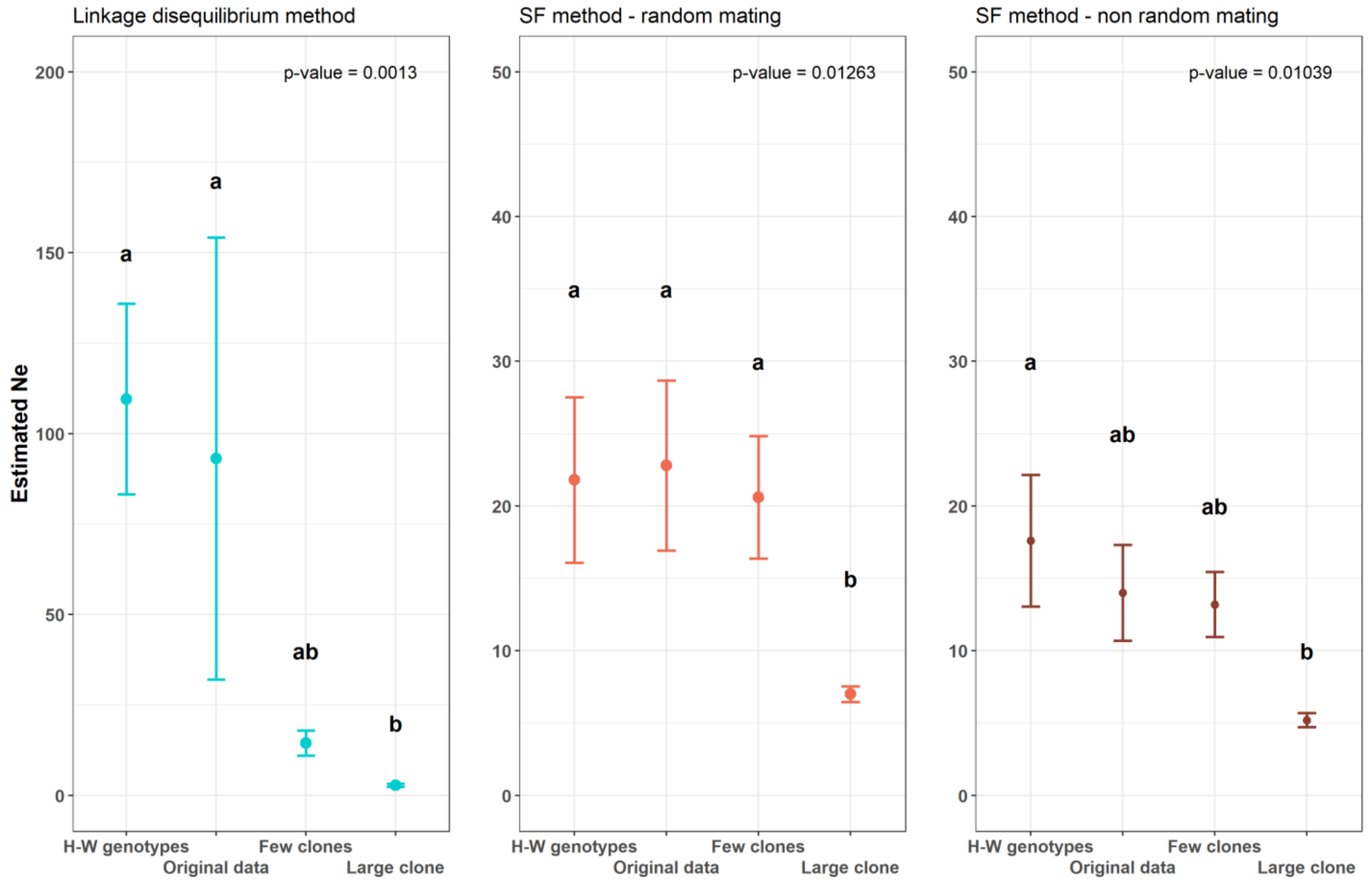

Figure S14. Mean values of $N_e$ estimates across an artificial clonality gradient and across five populations of *Suillus luteus*, *Rhizopogon togasawarius*, *Laccaria amethystina*, *Boletus edulis* and *Cantharellus formosus*. Each panel represents a different estimation method: Linkage disequilibrium, Sibship frequency (assuming random mating) and Sibship frequency (assuming non-random mating). Points represent the mean value across 5 populations, and error bars correspond to the standard error. Significant differences are indicated by letters and were tested using a Kruskal-Wallis rank sum test (p-value indicated in each panel), followed by a Dunn post hoc test (adjusted using the Holm method). Infinite values obtained using the LD method are not included in the graph, but were taken into account in the Kruskal-Wallis test.

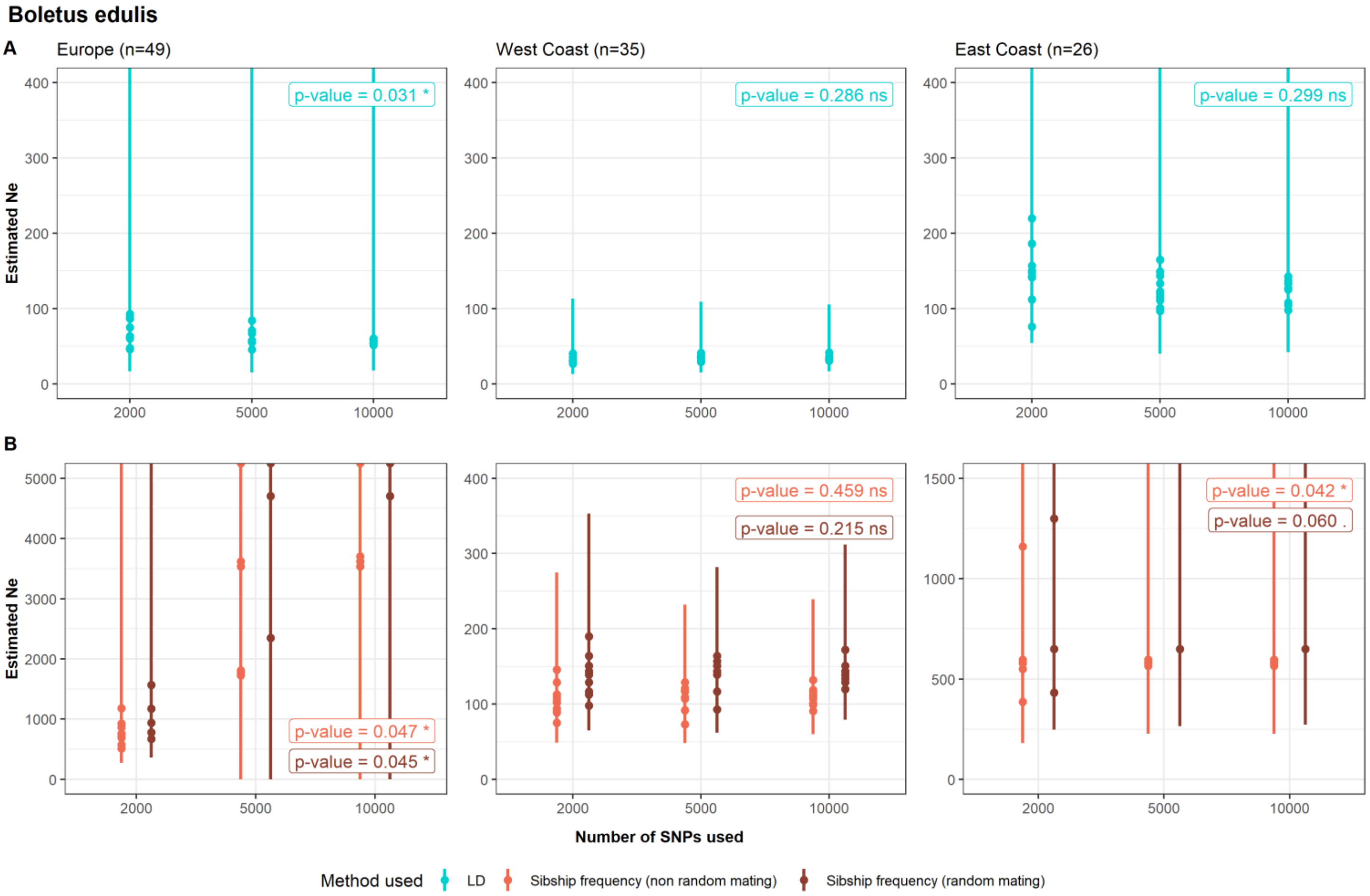

Figure S15. Estimation of $N_e$ with LD method (A) and SF method (B) in three populations of *Boletus edulis*. For each population, $N_e$ is estimated based on 10 subsets of 2,000, 5,000 and 10,000 SNPs from the original datasets. Each point represents the value of $N_e$ estimated by NeEstimator (LD) or Colony (SF), and the line range shows the confidence interval (Jackknife confidence interval for LD method, 95% confidence interval calculated from Student's distribution for the SF method). Points and lines that extend beyond the plot represent cases where the $N_e$ estimate or the upper limit of the confidence interval is infinite. P-values correspond to the results of a Levene's test to assess the homogeneity of variances of the $N_e$ estimates as a function of the number of SNPs used.

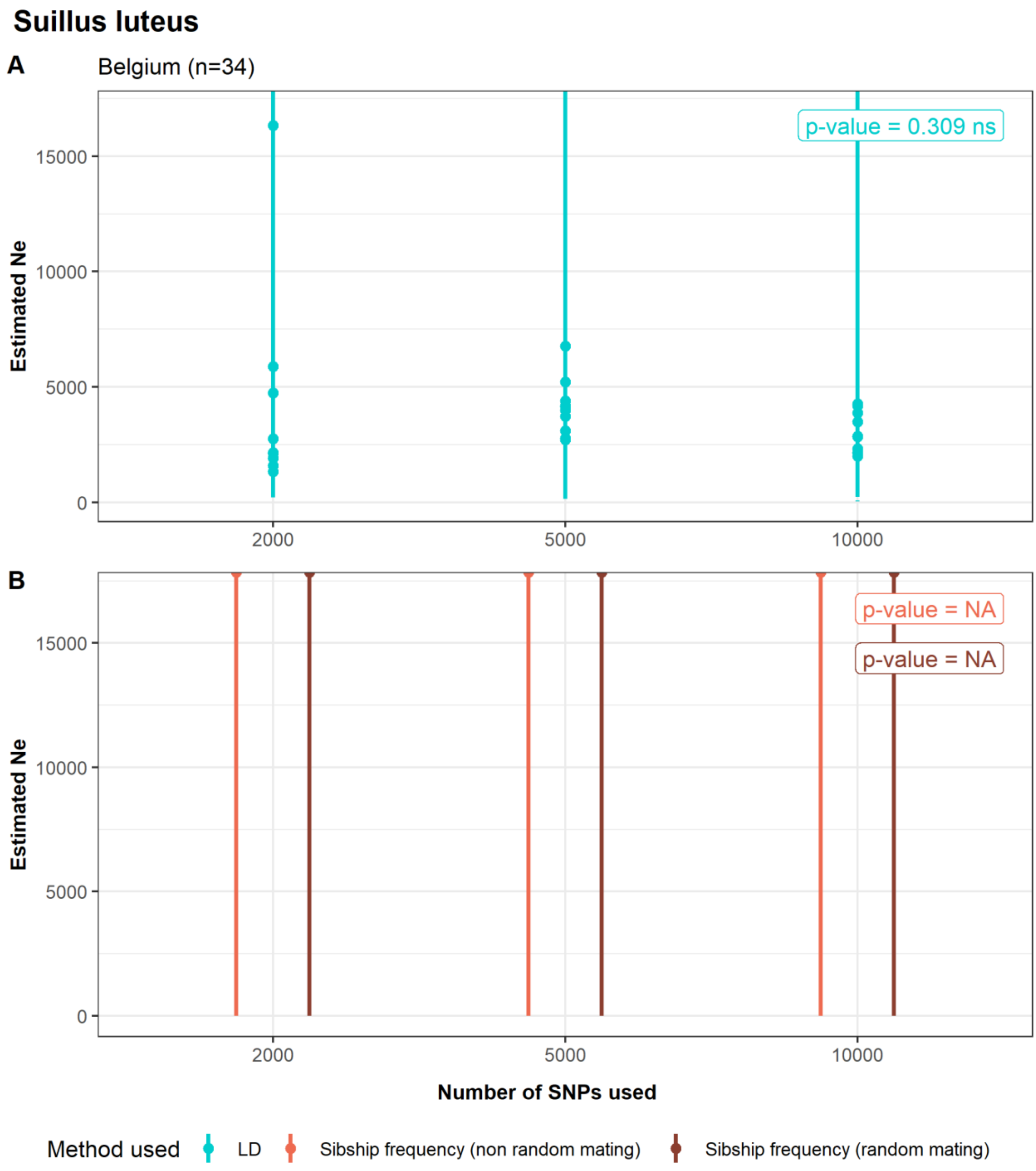

Figure S16. Estimation of $N_e$ with LD method (A) and SF method (B) in a population of *Suillus luteus*. For each population, $N_e$ is estimated based on 10 subsets of 2,000, 5,000 and 10,000 SNPs from the original datasets. Each point represents the value of $N_e$ estimated by NeEstimator (LD) or Colony (SF), and the line range shows the confidence interval (Jackknife confidence interval for LD method, 95% confidence interval calculated from Student's distribution for the SF method). Points and lines that extend beyond the plot represent cases where the $N_e$ estimate or the upper limit of the confidence interval is infinite. P-values correspond to the results of a Levene's test to assess the homogeneity of variances of the $N_e$ estimates as a function of the number of SNPs used.
